# Parallel evolution of plant alkaloid biosynthesis from bacterial-like decarboxylases

**DOI:** 10.1101/2024.06.04.597157

**Authors:** Catharine X. Wood, Zhouqian Jiang, Inesh Amarnath, Lachlan J. N. Waddell, Uma Sophia Batey, Oriana Serna Daza, Katherine Newling, Sally James, Gideon Grogan, William P. Unsworth, Benjamin R. Lichman

## Abstract

Alkaloids are nitrogen-containing natural products derived from amino acids. The basic amino acids lysine and ornithine are precursors to a wide range of alkaloids including the bioactive compounds nicotine, hyoscyamine and securinine. Feeding experiments have shown that the amino acids can be incorporated into alkaloids in a symmetric or nonsymmetric manner. The symmetric pathway is catalysed by two enzymes, a decarboxylase and oxidase, forming a cyclic iminium which acts as the electrophile in the scaffold forming step. Here, we describe the ornithine/lysine/arginine decarboxylase-oxidases (OLADOs), PLP-dependent enzymes responsible for the nonsymmetric pathway, catalysing the single step decarboxylative oxidative deamination of lysine, ornithine or arginine. These enzymes are group III ornithine/lysine/arginine decarboxylases (OLADs), an enzyme class previously exclusively associated with prokaryotes. We reveal OLADs to be widespread in plants and show that OLADOs have repeatedly emerged through parallel evolution from OLADs, *via* similar active site substitutions. This investigation introduces a new class of eukaryotic decarboxylases, and describes enzymes involved in multiple alkaloid biosynthesis pathways. It furthermore demonstrates how the principle of parallel evolution at a genomic and enzymatic level can be leveraged for gene discovery across multiple lineages.

## Introduction

Plant alkaloids are nitrogen-containing specialised metabolites derived from amino acids. They are notable for their complex polycyclic structures, significant bioactivities, and extensive roles as pharmaceuticals, components of traditional remedies and intoxicants. The basic amino acid lysine gives rise to various alkaloid classes across the green lineage, including piperidine alkaloids (anabasine, *Nicotiana sp*)^7^, *Lycopodium* alkaloids (huperzine A, *Huperzia serrata*)^8^, and quinolizidines (lupinine, *Lupinus sp*) (**Fig. 1A**)^9^. The non-proteogenic basic amino acid ornithine contributes to pyrrolidine alkaloids including nicotine and the tropane alkaloids (e.g. hyoscyamine) found in Solanceae species (**Fig. 1B**). Elucidating the biosynthesis of alkaloids can enable the formation of valuable pharmaceuticals in heterologous systems^10,11^ or the elimination of alkaloid biosynthesis in host tissues^9,12^.

**Figure 1.**
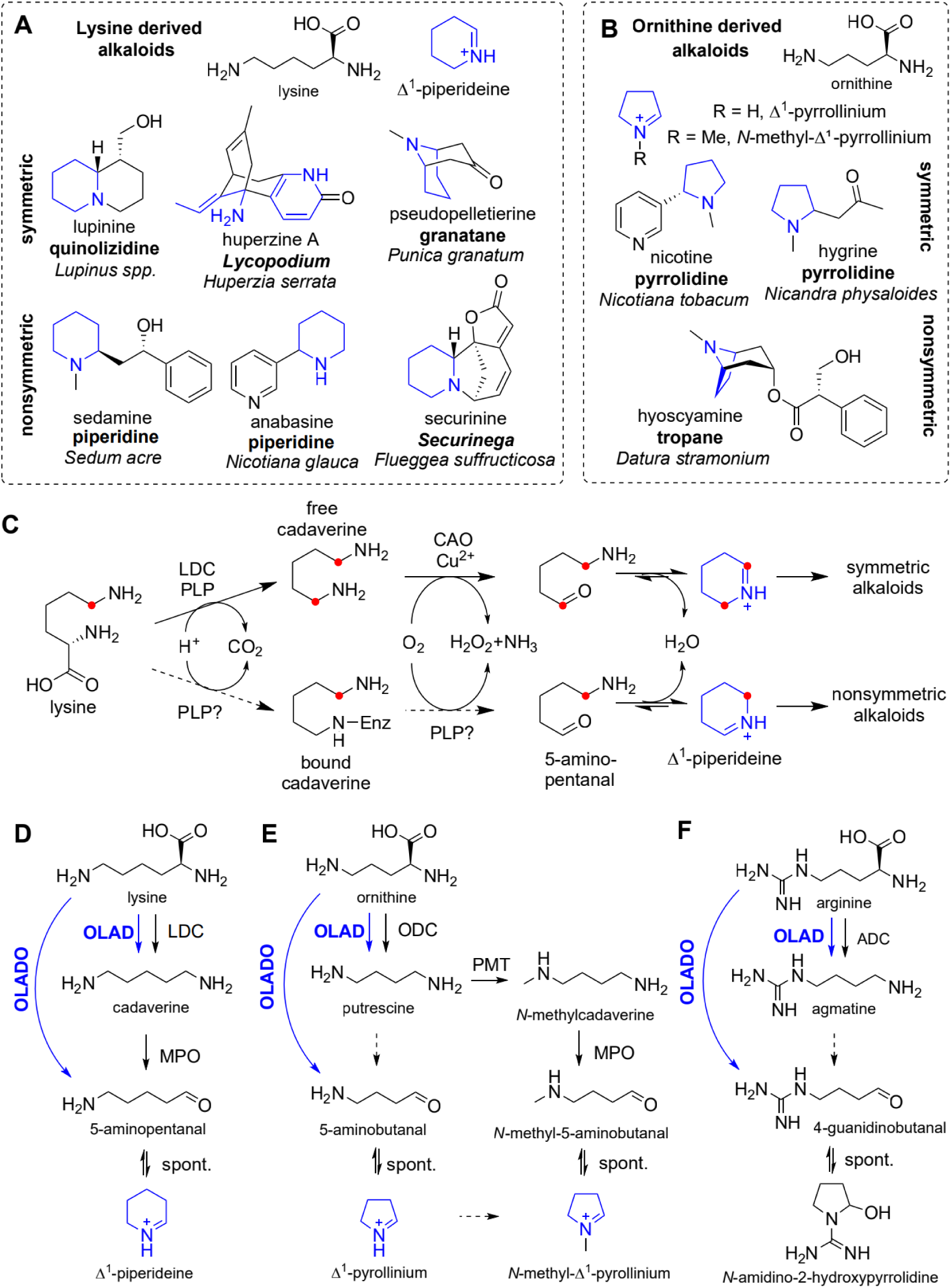
Basic amino acid derived alkaloids. **A.** Lysine and derived alkaloids formed *via* **Δ^1^**-piperideine (blue bonds). Alkaloid class in bold, example producing species in italics. **B.** Ornithine derived alkaloids formed *via* **Δ^1^**-pyrrollinium. **C.** Symmetric or nonsymmetric routes from lysine to Δ^1^-piperideine highlighting the fate of ε-carbon. The labels on the symmetric route Δ^1^-piperideine will be distributed on either carbon (but not both in the same molecule) whereas in the nonsymmetric route the label will be exclusively on the sp^3^ centre. The route from ornithine is analogous. **D-F.** Enzymatic decarboxylation and oxidation of basic amino acids lysine (**D**), ornithine (**E**) and arginine (**F**) with newly reported activities shown in blue.

Typical alkaloid biosynthesis pathways involve a scaffold-forming step wherein an iminium electrophile, derived from an amino acid, reacts with a nucleophile *via* a Mannich-like reaction^1^. For lysine and ornithine derived alkaloids the electrophiles are **Δ^1^**-piperideine or **Δ^1^**-pyrrollinium (or methylated derivatives) which form from the amino acids *via* decarboxylation, oxidation and cyclisation (**Fig. 1C**). The feeding of isotopically labelled precursors to plants revealed that the formation of **Δ^1^**-piperideine or **Δ^1^**-pyrrollinium could occur either in a symmetric or nonsymmetric manner, with the former indicating the presence of a free intermediate (e.g. cadaverine), whereas the latter was proposed to proceed *via* an enzyme bound intermediate^13^.

The enzymatic origins of symmetric Δ^1^-piperideine from the *Lycopodium*^4^ and quinolizidine^14^ alkaloid families have been determined, and involve two enzymes: a PLP-dependent lysine decarboxylase (LDC) catalysing the formation of cadaverine from lysine, followed by a copper amine oxidase (CAO) catalysing the oxidation of cadaverine into 5-aminopentanal, which spontaneously cyclises into **Δ^1^**-piperideine (**Fig. 1C** and **D**). Analogous pathways from ornithine to symmetric pyrrolidine or tropane alkaloids in Solanaceae including *Atropa belladonna*^15^ and *Nicotiana*^16^ have been proposed, *via* putrescine, though include a methylation step prior to oxidation (**Fig. 1E**)^17^.

Piperidine alkaloids anabasine^18^ and sedamine (*Sedum acre*)^13^ are known to have nonsymmetric origins, as does the *Securinega* alkaloid securinine, a GABA-receptor antagonist from *Flueggea suffruticosa* (**Fig. 1A**)^19^. There is also evidence of nonsymmetric labelling in tropane alkaloids from *Datura stramonium*^20^. In 1973, Leistner and Spenser hypothesised that a PLP-dependent oxidative enzyme could be responsible for the nonsymmetric pathway to Δ^1^-piperideine^13^, but to date no enzymes responsible for nonsymmetric alkaloid biosynthesis have been described.

Parallel evolution, a subset of convergent evolution in which new activities arise independently from within the same enzyme family, is a common phenomenon in plant specialised metabolism, including alkaloid biosynthesis^21^. For example, piperidine alkaloid related LDCs have evolved multiple times independently from ornithine decarboxylases (ODCs), members of the group IV PLP-dependent decarboxylase family^4^. Homospermidine synthase, a key enzyme in pyrrolizidine alkaloid biosynthesis, has evolved on multiple occasions through duplication of the essential enzyme deoxyhypusine synthase^22^. If one enzyme responsible for nonsymmetric Δ^1^-piperideine or **Δ^1^**-pyrrollinium formation were to be characterised, it may be possible to use the principle of parallel evolution to identify functionally equivalent enzymes from other lineages.

Our investigation began with the goal of identifying the early steps of securinine biosynthesis in *F. suffruticosa*, as part of a broader effort to elucidate the pathway of the *Securinega* alkaloids^23^. In doing so, we discovered an enzyme (Fs1864) capable of catalysing the nonsymmetric oxidative formation of **Δ^1^**-piperideine from lysine (**Fig. 1D**). This enzyme is a member of the PLP-dependent group III ornithine/lysine/arginine decarboxylase (OLAD) family, which is typically associated with prokaryotes. We showed that Fs1864 has ornithine/lysine/arginine decarboxylating deaminative oxidase (OLADO) activity whereas a paralog (Fs3498) catalyses decarboxylation without oxidation (**Fig. 1D, E, F**). Using mutagenesis, we identify key amino acids contributing to the substrate acceptance and oxidation activity. The expectation that other nonsymmetric alkaloid pathways emerged *via* parallel evolution of OLADO enzymes led to the identification and characterisation of equivalent pairs of OLAD/OLADO enzymes in the alkaloid producing *Nicotiana tabacum* and in *Artemisia annua*, which is not yet known to produce alkaloids.

This work resolves the longstanding question of nonsymmetric piperidine alkaloid biosynthesis, and in doing so also (i) describes key enzymes in securinine and anabasine biosynthesis, (ii) reveals a new class of eukaryotic bacterial-like group III decarboxylases, (iii) demonstrates novel OLADO activities, and (iv) highlights how the expectation of parallel evolution can be used for gene discovery.

## Results

### Identification of Δ^1^-piperideine forming enzyme

We set out to discover a lysine decarboxylase (LDC) from *F. suffruticosa* that could catalyse the first step in the formation of **Δ^1^**-piperideine that contributes the piperidine moeity in securinine. First, we generated a *F. suffruticosa* transcriptomic resource from 15 different tissues and developmental stages (**Table S1**), combining short and long-read sequencing to assemble a *de novo* transcriptome and gene expression information (**Fig. S1**). Using functional annotation, we identified five gene candidates identified as members of ornithine decarboxylase families (**Fig. 2A**). This term was selected as plant LDCs are known to be closely related to ornithine decarboxylase^4^. Two candidates were filtered out due to apparent transcriptome misassembly (gene 25968, **Fig. S2**) and low gene expression (gene 3498, **Fig. S3**).

**Figure 2.**
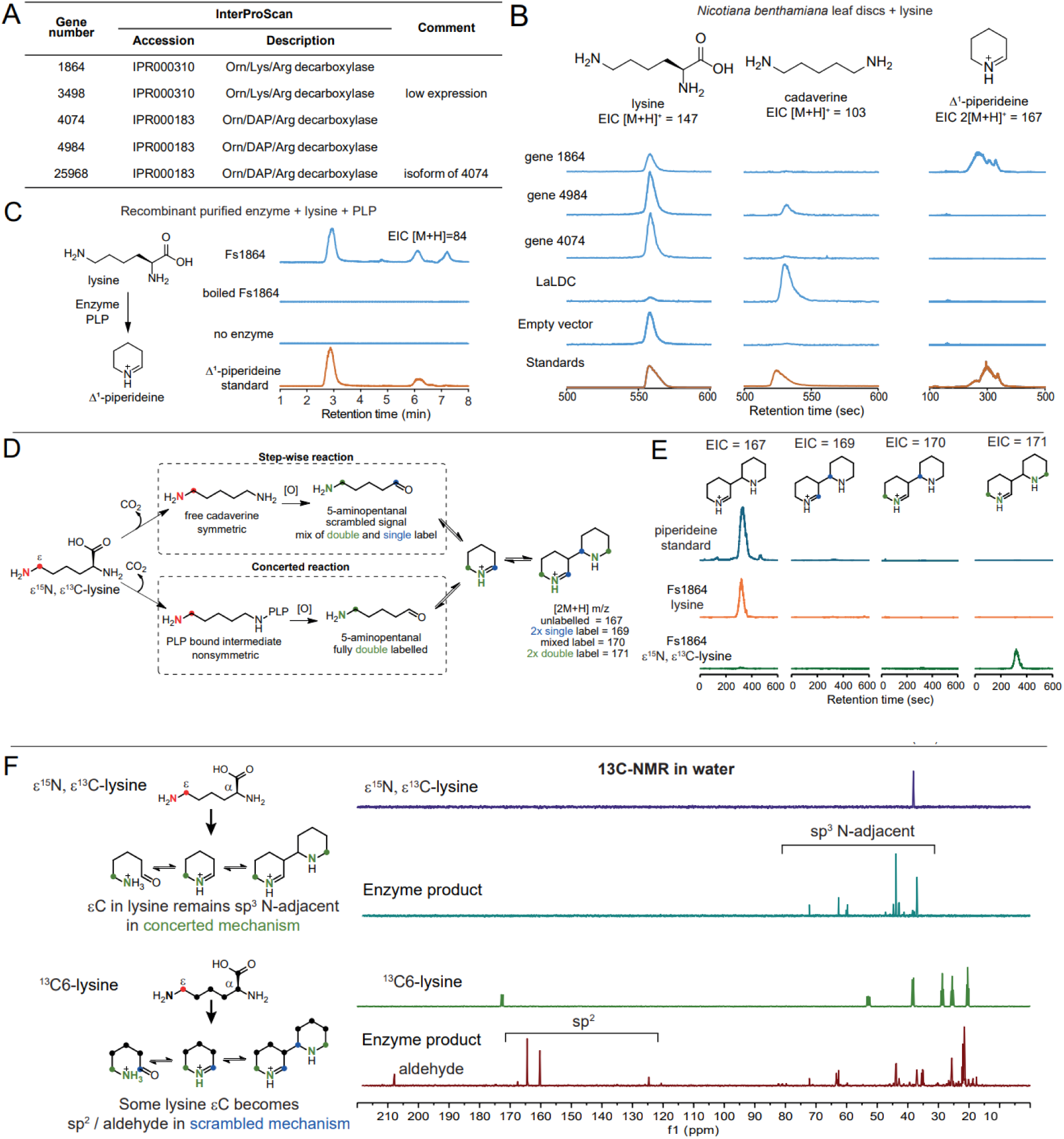
Identification of Fs1864 catalysed non-symmetric Δ^1^-piperidiene formation. **A.** LDC candidates. **B.** LC-MS analysis of leaf disc assays. Representative extracted ion chromatograms (EIC) for (L-R) lysine, cadaverine and Δ^1^-piperideine dimer. Peak height in each EIC is comparable across samples; standard intensities were scaled. *Lupinus angustifolius* LDC (LaLDC) is positive control, empty vector (EV) is negative control. **C.** *In vitro* assay of recombinant Fs1864 (20 ng/µL) showing formation of Δ^1^-piperidiene from lysine (10 mM) with additional PLP. **D.** Isotope labelling outcomes of stepwise (i.e. free cadaverine intermediate) or concerted (i.e. bound intermediate) formation of Δ^1^-piperideine using ε^15^N, ε^13^C-lysine. **E.** LC-MS results of Fs1864 reaction with unlabelled lysine or ε^15^N, ε^13^C-lysine. EICs of Δ^1^-piperideine dimers: unlabelled (167), 2x single labelled (169), single+double labelled (170) and 2x double labelled (171). **F.** ^13^C-NMR spectra of ε^15^N, ε^13^C-lysine substrate and Fs1864 product compared to fully labelled 13C6-lysine and Fs1864 product.

We cloned the three remaining candidates from *F. suffruticosa* (**Table S2**) and, using *Nicotiana benthamiana* transient expression leaf disc assays, we tested their activity with lysine. Using LC-MS, we observed cadaverine formation in the positive control (*Lupinus angustifolius* LDC)^4^ and gene 4984 (**Fig. 2B, Fig. S4**). For gene 1864, we saw consumption of lysine, no accumulation of cadaverine, but accumulation of Δ^1^-piperideine, suggesting direct conversion of lysine to Δ^1^-piperideine.

### *In vitro* validation of non-symmetric Δ^1^-piperidiene formation from lysine

To verify the activity of the enzyme encoded by *F. suffruticosa* gene 1864 (Fs1864), we expressed it recombinantly in *E. coli* as an Δ74N-terminal truncate, based on AlphaFold structural model analysis (**Fig. S5A, Table S3**). Incubation of the purified enzyme (**Fig. S5B**) with lysine and PLP led to the formation of Δ^1^-piperideine (**Fig. 2C**). Coupling the reaction to a peroxidase assay showed H_2_O_2_ accumulation over the course of the reaction (**Fig. S5C**), which is enhanced by supplemental PLP co-factor (**Fig. S5D**). This supports Fs1864 producing Δ^1^-piperideine from lysine *via* a PLP-dependent decarboxylation-oxidative deamination reaction^24^.

Isotope labelling experiments in *F. suffruticosa* had shown that Δ^1^-piperideine in securinine originates from lysine in a nonsymmetric manner^25^. Having identified Fs1864 as the candidate enzyme for this process, we were able to examine this phenomenon *in vitro*. There are two possible routes to Δ^1^-piperideine from lysine (**Fig. 2D**). In the step-wise route, where the reaction occurs *via* a free cadaverine intermediate, we would expect to have a scrambled, symmetric signal and to observe both singly and doubly labelled products, which would provide a mixture of three dimer masses, [2M+H]^+^ = 169, 170 and 171. In a concerted route, where the decarboxylated intermediate is enzyme bound, we would observe only the double labelled product: [2M+H]^+^ = 171. When Fs1864 was supplied with ε^15^N, ε^13^C-lysine, only the doubly labelled product was observed, indicating a concerted reaction (**Fig. 2E**).

To determine the position of the isotope labels on the product, we analysed the enzyme product using ^13^C-NMR (**Fig. 2F**). The spectra were obtained directly from enzyme assays conducted in water; hence the product is present as a mixture of forms. With ε^15^N, ε^13^C-lysine as the substrate, we observe ^13^C-NMR peaks at 30-70 ppm, consistent with sp^3^-hybridrised carbon atoms adjacent to a heteroatom. As a control, we compared the peaks to the full ^13^C-NMR spectrum of the enzyme product in water, derived from the fully ^13^C-labelled ^13^C_6_-lysine. In the product, we noted the presence of sp^2^ hybridised (likely imine/iminium signals) and aldehyde peaks, which may be derived from the lysine αC or εC if the reaction occurred *via* a free cadaverine, but only the αC in a concerted reaction. These peaks are absent in the ε^15^N, ε^13^C-lysine derived reaction, suggesting the equivalent sp^2^ and aldehyde peaks are derived from the unlabelled αC and therefore not visible on the spectrum.

This *in vitro* labelling study confirms, using both mass-spectrometry and ^13^C-NMR, a concerted reaction mechanism that proceeds without a free intermediate and provides an enzymatic explanation for the nonsymmetric labelling observed in the plant.

### Ornithine/lysine/arginine decarboxylases

LDCs are not known to be universally found in plants: characterised LDCs operating in plant specialised metabolism have evolved from ODCs on multiple independent occasions^20^. The tendency of plant LDCs to evolve convergently alongside the unprecedented nonsymmetric oxidative deamination of lysine catalysed by Fs1864 prompted an investigation into the enzyme’s evolutionary origin.

Based on annotation and sequence similarity, Fs1864 was revealed to be a member of the PLP-dependent group III decarboxylase family (InterPro: major domain, IPR000310; family IPR011193)^5,26^. This group is part of the larger PLP-dependent fold I superfamily, which also includes group II decarboxylases (InterPro: family IPR002129), which feature plant amino acid decarboxylases and aldehyde synthases^27,28^. Notably this is a different fold type and decarboxylase group to typical plant LDC/ODCs, which are group IV decarboxylases (InterPro: family IPR000183), with PLP fold type III. Structures of representatives of these sequences including Fs1864 confirm the sequence-based relationships (**Fig. 3A**). The decarboxylase group III is known as the prokaryotic type ornithine/lysine/arginine (Orn/Lys/Arg) decarboxylases (OLADs)^5^. To our knowledge, there is only a single report mentioning eukaryotic OLADs^29^, and none has been characterised.

**Figure 3.**
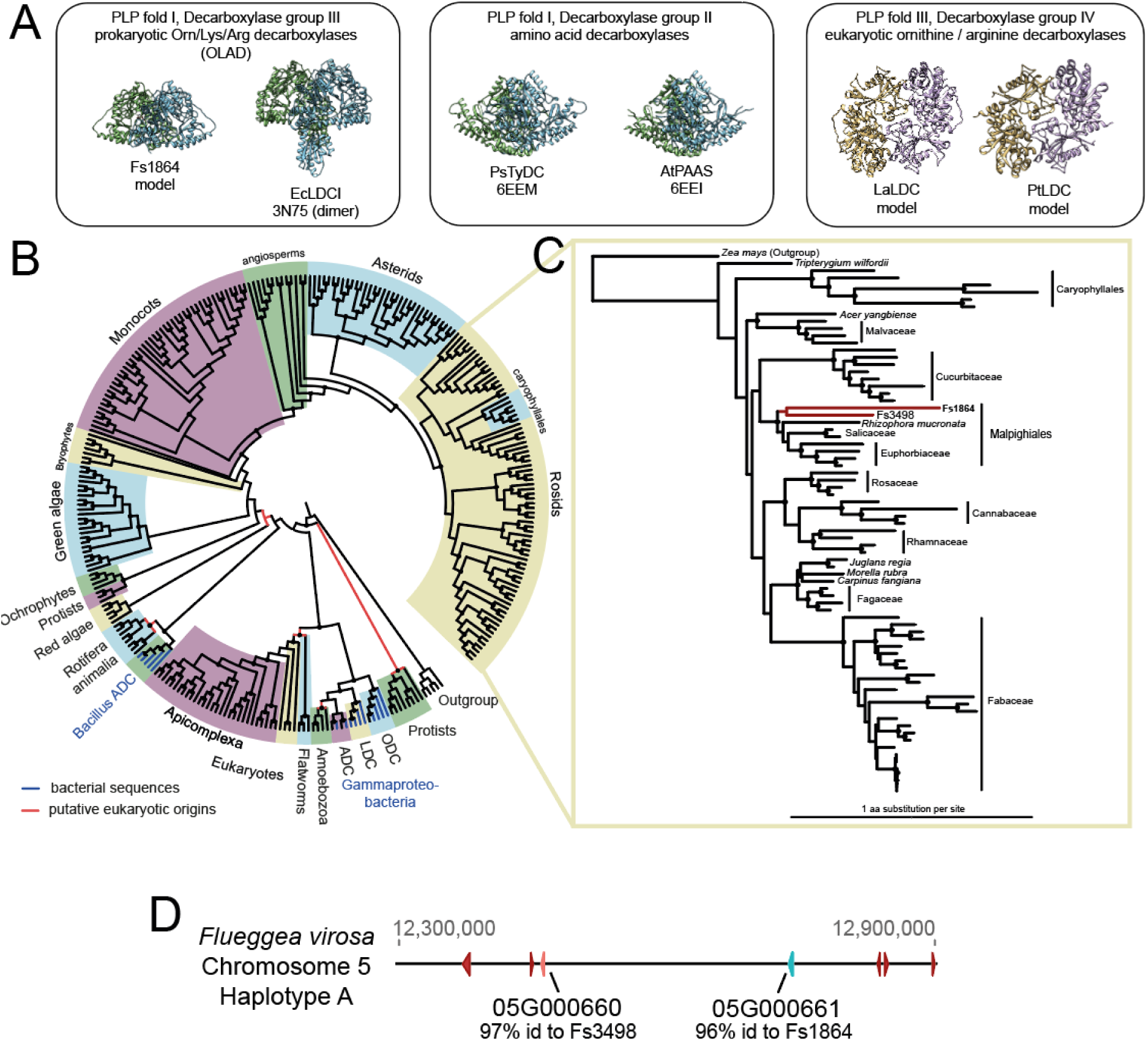
Evolutionary origins of Fs1864. **A.** X-ray structures or structural models of PLP-dependent decarboxylases. **B.** Maximum likelihood phylogeny of selected group III decarboxylase protein sequences. Taxonomic clades highlighted by colours. Sequences derived from bacteria are in blue, all other sequences are uncharacterised eukaryotic sequences. Red branches show putative eukaryotic origins. Black dots show highly supported nodes (SH-aLRT ≥80% and UFboot ≥95%). Sequence names and branch lengths on **Fig. S6**. **C.** Focussed rosid phylogeny of selected group III decarboxylase protein sequences. Fs1864 and Fs3498 highlighted in red. **D.** Genomic location of group III decarboxylases in *Flueggea virosa* genome, showing signature of tandem duplication. Amino acid identities are based on the predicted structured regions (as determined by AlphaFold).

We constructed a phylogenetic tree of OLADs, based on the presence of the major domain (InterPro: IPR000310) and focussing on eukaryotic sequences (**Fig. 3B, Fig. S6**). Ignoring single reports from a clade, which could be the result of sample contamination, the phylogenetic tree reveals at least four origins of eukaryotic OLADs. There appear to be gammaproteobacteria arginine decarboxylase (ADC) like genes in Amoebozoa. There is also a eukaryotic lineage sister to gammaproteobacterial OLADs found in flatworms and Apicomplexa protists. Bacillus-like ADCs appear in rotifers, microscopic animals.

The eukaryotic OLAD lineage leading to FsOLADO has an algal origin and is found in red, green and ochrophyte algae. It is also widespread in green plants: there are orthologs in bryophytes, monocots (including rice, maize and wheat), asterids and rosids. However, no homologs were identified in gymnosperms nor in Arabidopsis. Examination into the origin of Fs1864 appears to have revealed a hitherto neglected family of plant enzymes, the OLADs, which may have wider roles in polyamine biosynthesis.

### F. suffruticosa OLAD and OLADO

To examine Fs1864 evolution more closely, we constructed a phylogenetic tree of OLADs from the rosid clade of flowering plants (**Fig. 3C**). This analysis revealed the presence of homolog of Fs1864 also present in *F. suffruticosa.* This paralog is Fs3498, which was previously excluded from screening due to low expression. Fs1864 and Fs3498 are sister sequences yet share only 54% sequence identity. Furthermore, Fs1864 has <60% sequence identity with orthologs from Malpighiales, whereas Fs3498 has >65% identity with the same sequences. Altogether this indicates a gene duplication and activity divergence event in the *Flueggea* lineage, where Fs3498 maintained ancestral-like OLAD activity and Fs1864 evolved new oxidative activity related to alkaloid biosynthesis. Furthermore, we examined the genome assembly of *F. virosa*, which also produces securinine-type alkaloids. No genome is currently available for *F. suffruticosa*. We identified orthologs of Fs1864 and Fs3498 present as tandem duplicates on *F. virosa* chromosome 5 (05G000661 and 05G000660 respectively), a genomic signature of lineage specific neofunctionalisation (**Fig. 3D**)^30^.

Based on this hypothesis, we elected to test the activity of Fs1864 and Fs3498 on lysine, ornithine and arginine. *In vitro*, all three amino acids were substrates for the enzymes, with Fs3498 performing decarboxylation and only trace oxidation, whereas Fs1864 was able to perform both decarboxylation and oxidative deamination on all three substrates (**Fig. 4A**). We consequently renamed Fs3498 to FsOLAD and Fs1864 to FsOLADO. The ability of FsOLADO to make both Δ^1^-piperideine and Δ^1^-pyrrollinium is notable as this mirrors the presence in *F. suffruticosa* of securinine and its pyrrolidine analog norsecurinine. FsOLADO is also the first reported plant enzyme catalysing formation of γ-guanidinobutyraldehyde (GBAL) from arginine.

**Figure 4.**
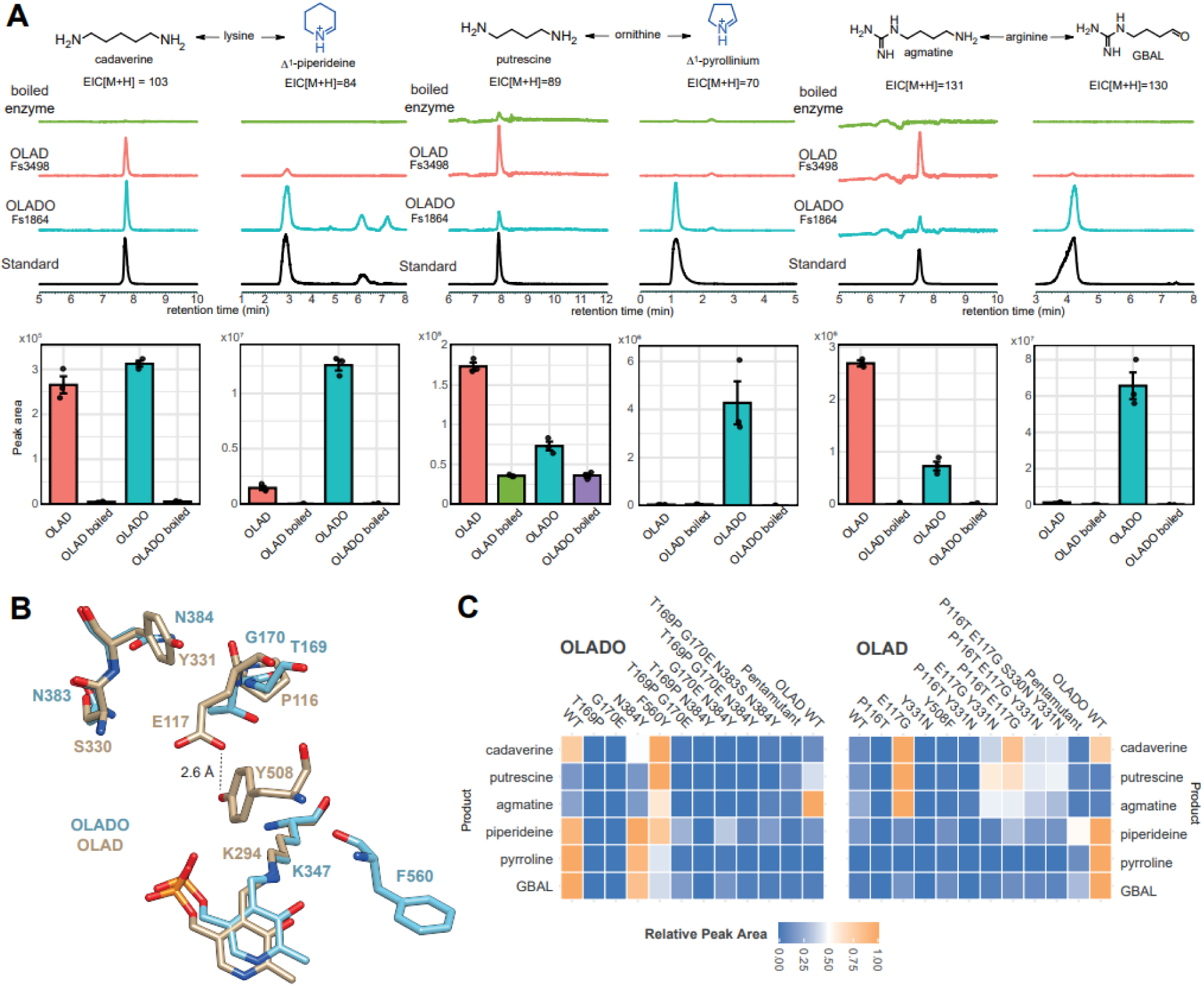
Activities of FsOLAD and FsOLADO. **A.** Activity of purified enzymes with lysine, ornithine and arginine. LC-MS analysis of *in vitro* assays, forming either the decarboxylated product (cadaverine, putrescine or agmatine) or decarboxylated and oxidised (and cyclised) product (Δ^1^-piperideine, Δ^1^-pyrrollinium and γ-guanidinobutyraldehyde [GBAL]). Representative extracted ion chromatograms (EIC) depicted, where peak height within each EIC is comparable across samples. Negative control is boiled FsOLAD. Bar graphs show corresponding EIC peak areas; bars show mean of triplicate reactions, points from each reaction (n = 3), error bars are SE. Substrate concentration 10 mM, enzyme concentration 20 ng/µL. **B.** Active site comparison of FsOLAD (beige) and FsOLADO (blue) showing PLP-binding lysine alongside residues proposed to influence activity. **C.** Heat map showing *in vitro* product formation in purified enzyme variants, with variants shown in columns and products shown in rows, with colours relative to maximum reported conversions in each row. Pentamutants have all residues (as depicted in panel B) swapped. Substrate concentration 5 mM, enzyme concentration 20 ng/µL. Values are the mean of at least three assays, showing in full in **Fig. S9**.

### Active site variants

We aimed to understand the structural basis of the different activities of FsOLAD and FsOLADO. To do this, we compared protein models generated by AlphaFold and inspected the active site, alongside sequence alignments. Three regions of interest were identified where residues FsOLADO appeared to provide greater space in the active site than those in FsOLAD: N383-N384 (OLADO) compared to S330-Y331 (OLAD), T169-G170 compared to P116-E117, and F560 compared to Y508 (**Fig. 4A** and **Fig. S7**). In FsOLAD, E117 and Y508 side-chains interact through an H-bond. This interaction is absent in FsOLADO which appears to result in a notable increase in active site volume. The N384/Y331 position aligns with the residue found to be involved in the transition between decarboxylation (tyrosine) and aldehyde synthase (phenylalanine) activities in the distantly related group II decarboxylases (**Fig. S8A**)^31^.

To examine the role of these residues we substituted them to the corresponding residue in the paralog (**Table S3** and **S4**), purified the enzymes and tested the variants on the three substrates (**Fig. 4C** and **Fig. S9**). For FsOLADO, T169P and G170E generally appeared disruptive to both decarboxylation and oxidation activities. N384Y was largely undisruptive to decarboxylation but appeared to boost oxidase activities, especially in combination with other substitutions. The F560Y substitution increased decarboxylation activity but reduced oxidation. For FsOLAD, E117G boosted decarboxylation activity, both alone and in combination with other substitutions. The substitution most influential for oxidation activity was Y508F, especially when introduced to the P116T/E117G/S330N/Y331N variant where it both reduced decarboxylation and increased oxidation. Overall, across FsOLAD and FsOLADO, the key residue for determining decarboxylation or oxidation activity appears to be Y508/F560.

Based on these results, we speculate that in FsOLAD Y508 donates a proton to the intermediate-PLP complex to trigger imine formation and the decarboxylation product (**Fig. S10**). Therefore in FsOLADO F560Y decarboxylation is enhanced. We also propose that in FsOLAD E117 is H-bonded to Y508 and restricts the active site volume, preventing effective substrate binding: the OLAD-E117G substitution enables improved substrate binding and boosts decarboxylation. The impact of Y508F on OLAD oxidation is only clear in the presence of E117G as the substrate is required to bind. This interpretation also implies that the natural substrate of FsOLAD may not be lysine, ornithine or arginine but a smaller molecule.

### Parallel evolution of OLADO from OLAD in Solanaceae

Convergent evolution of enzyme activity or gene function is not a rare phenomenon in plant specialised metabolism^6^. In particular, parallel evolution, which is the independent emergence of activity in separate lineages but from similar ancestors (i.e. same enzyme class) appears to be commonplace^6,32^. Two relevant examples are the parallel evolution of canonical LDCs from ODCs^4^, and the development of aldehyde synthase activities from group II decarboxylases^31^. Parallel evolution may even involve convergence of amino acid sequences, especially for active site residues^32^. We predicted that multiple lineage specific nonsymmetric alkaloid biosynthesis pathways were controlled by OLADO enzymes that had evolved, in parallel, from the OLAD family. As in *Nicotiana* sp. Δ^1^-piperideine is incorporated into anabasine in a non-symmetric manner (**Fig. 1A**)^18^, we predicted an OLADO would be responsible for Δ^1^-piperideine formation in these species. To investigate this, we inferred a focussed phylogeny of OLADs from the asterid clade, and examined the active site residues for similarities to FsOLADO. This investigation revealed two clades in the asterids showing OLADO-like residues, notably phenylalanines in the FsOLADO-F560 position and no tyrosine in the FsOLADO-N384 position (**Fig. 5**). One of these OLADO-like clades featured solely Solanaceae species, an alkaloid-rich family which includes *Nicotiana* and tropane alkaloid producing species such as *Datura stamonium*. Another clade with OLADO features contained Asteraceae species including *Artemisia annua*, which is not known to produce alkaloids. For both taxa (Solanaceae and Asteraceae) there were also clades that lack OLADO sequence features, which were presumably OLAD paralogs.

**Figure 5.**
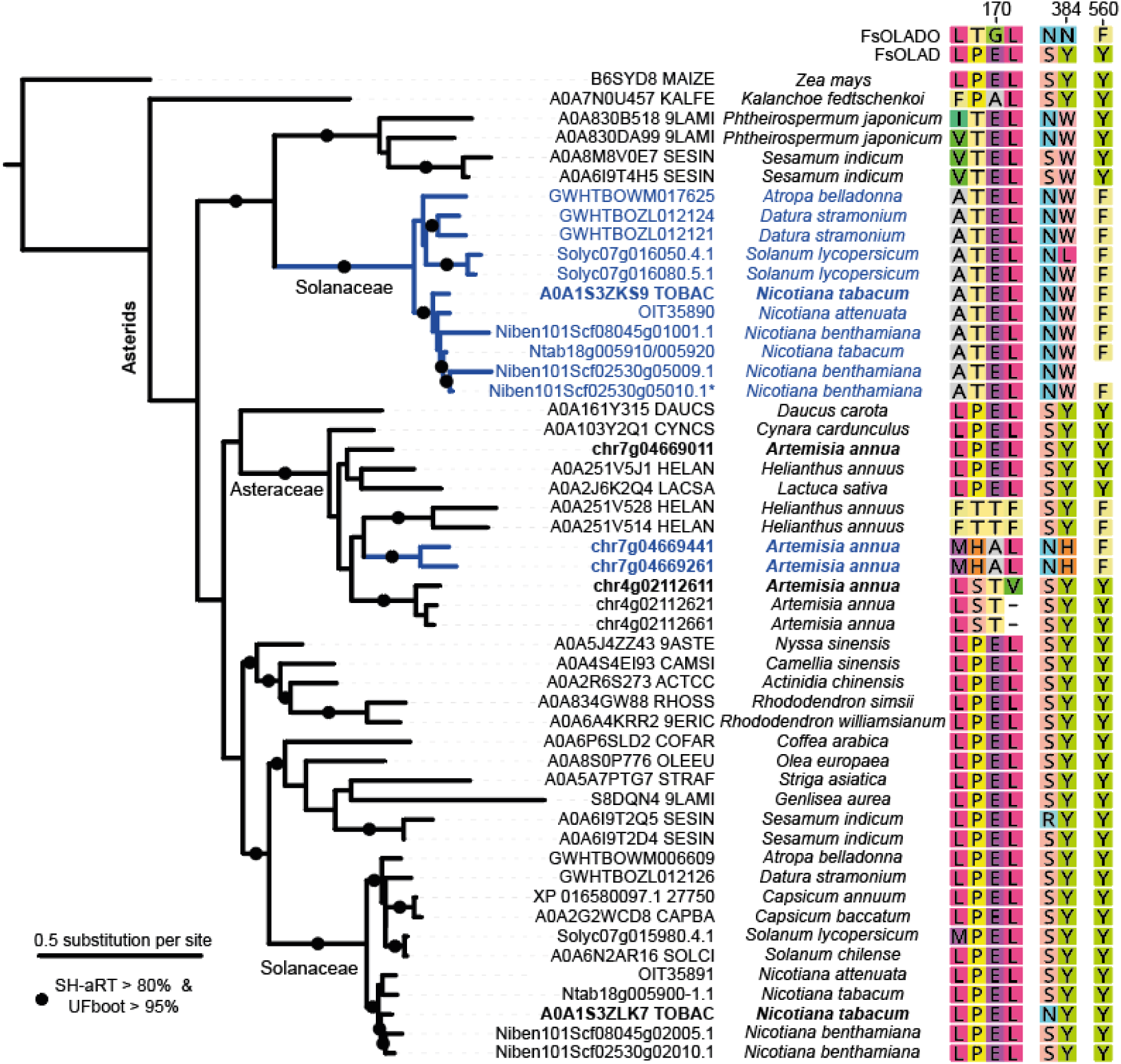
Parallel evolution of OLADO in Asterids. Focussed Asterid phylogeny of selected group III decarboxylase (OLAD) protein sequences. Black dots show highly supported nodes (SH-aLRT ≥80% and UFboot ≥95%). Bold sequences are those that were selected for testing (Fig. 6). Blue clades are OLADO-like due to key residue identities. Selected alignment columns show identity of key residues in selected sequences compared to FsOLAD/OLADO. Residue numbering is based on FsOLADO. Starred *N. benthamiana* OLADO sequence was used for co-expression analysis (**Fig. S12**, **Table S5**).

We elected to investigate OLAD/OLADO activities in *N. tobacum*, which produces anabasine as a minor alkaloid component^7^. Closer investigation into the *Nicotiana tobacum* genome assembly revealed five OLAD paralogs localised across two chromosomes (**Fig. 6A**), though the published gene models failed to accurately define the gene boundaries so these were refined by cross reference with other datasets (**Fig. S11**). Each region contains tandem genes from the two distinct Solanaceae OLAD clades (**Fig. 5**), suggesting origins *via* a combination of tandem gene duplication and whole genome duplication. Due to the high similarity between the paralogs within each clade, we elected to test the activity of one representative from each clade, both from chromosome nine: A0A1S3ZLK7 from the OLAD clade and A0A1S3ZKS9 from the proposed OLADO clade (**Table S3**). As hypothesised, the activities closely resembled the FsOLAD/FsOLADO pair, with A0A1S3ZLK7 demonstrating decarboxylation of lysine, ornithine and arginine but minimal oxidation and A0A1S3ZKS9 showing oxidative deamination of lysine, ornithine and arginine, alongside the decarboxylation (**Fig. 6B**). Based on this activity we renamed the enzymes NtOLAD and NtOLADO respectively. We would expect other enzymes (paralogs or orthologs) in the same Solanaceae clades and sharing the key active site residues, to share similar distinct OLAD and OLADO activities (**Fig. 5**).

**Figure 6.**
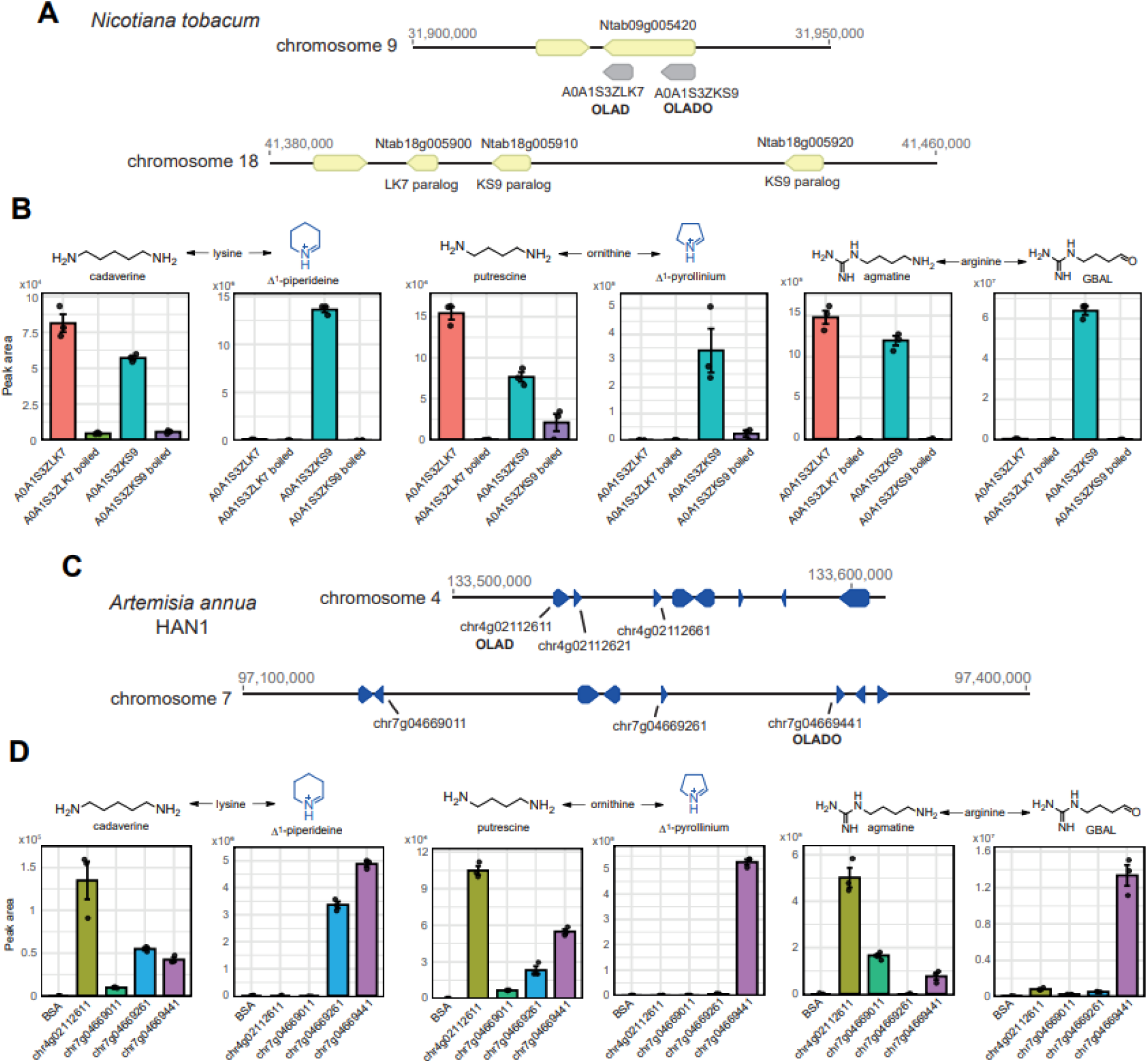
Activities of Asterid OLAD/OLADOs. **A.** Genomic location of *N. tabacum* OLAD paralogs. Refined gene model of Ntab09g005420 shown in **Fig. S11**. **B.** Activity of purified enzymes with lysine, ornithine and arginine. Bar graphs show LC-MS derived EIC peak areas; bars show mean of triplicate reactions, points from each reaction (n = 3), error bars are SE. **C.** Genomic location of *A. annua* OLAD paralogs. **D.** Activity of purified enzymes with lysine, ornithine and arginine. Bar graphs show LC-MS derived EIC peak areas; bars show mean of triplicate reactions, points from each reaction (n = 3), error bars are SE, substrate concentration 10 mM, enzyme concentration 10 ng/µL.

Gene expression patterns would provide further insight into the role of these sequences. Due to the inaccurate gene models we were unable to assess *N. tobacum* expression using existing resources but instead used *N. benthamiana* data, which also produces nicotine and anabasine^7,33^. The two *N. benthamiana* OLAD genes showed balanced expression across multiple tissues, contrasting with OLADOs, which were more varied across tissues and included pseudogenes with near zero expression (**Fig. S12**). In *N. benthamiana*, two OLADO paralogs (Scf02530g05010 and Ctg12894g00002) had high expression in root and stem samples. These genes had very high expression correlation with orthologs from the nicotine biosynthesis pathway (top 30 correlating genes, PCC = 0.75–0.93, **Table S5**). This assessment suggests that OLADOs are involved in *Nicotiana* alkaloid biosynthesis.

### Parallel evolution of Asteraceae OLADO

Through our examination of OLAD genes in the asterid clade, we noticed a clade in of Asteraceae sequences that had distinctly OLADO-like residues: phenylalanines in the FsOLADO-F560 position and no tyrosine in the FsOLADO-N384 position (**Fig. 5**). We elected to examine these enzymes further, focussing on *Artemisia annua*, despite there being limited reports of alkaloids in this species^34^. Analysis of the genome assembly revealed six genes across two chromosomes, in arrays of three. Based on the phylogeny, these appear to be the outcome of Asteraceae-specific tandem and whole genome duplications (**Fig. 5**). We expressed, purified and characterised four of these enzymes, two hypothesised to be OLAD and two OLADO based on active site residues (**Fig. 5, Table S3**). The hypothesised OLADs chr7g04669011 and chr4g02112611 indeed showed decarboxylation activity and minimal oxidation activity with lysine, ornithine and arginine, though notably chr4g02112611 had superior activity across all substrates. This may be because it has a threonine present in the FsOLADO-E383 position (**Fig. 5**), which based on the mutation analysis (**Fig. 4**) would open up space in the active site for these substrates. The predicted OLADOs demonstrated decarboxylation and oxidation activities, with chr7g04669441 turning over all three substrates tested but chr7g04669261 only showing comparable activity on lysine.

## Discussion

The nonsymmetric incorporation of basic amino acids into alkaloids was noted in early biosynthetic investigations employing radioactively labelled precursors^35^. Later, Leistner and Spenser proposed that a PLP-dependent oxidase could be responsible for the direct nonsymmetric conversion of lysine into Δ^1^-piperideine^13^. Despite recent advances in elucidating the elaborate pathways of alkaloid biosynthesis^8,36^, the nature of this crucial early step has been unresolved until now.

In this work we have revealed the plant enzymes that are capable of the nonsymmetric formation of lysine and ornithine derived alkaloids: the OLADOs. We first identified an OLADO in *F. suffruticosa*, showing it to be a PLP-dependent enzyme capable of producing Δ^1^-piperideine from lysine *via* decarboxylation and oxidation, validating the previous hypothesis^13^. In the putative *Securinega* alkaloid pathway, OLADO’s products Δ^1^-piperideine and Δ^1^-pyrrolinium are precursors to alkaloids such as securinine and norsecurinine. This work therefore describes the first enzyme of *Securinega* alkaloid biosynthesis, enabling future elucidation of the full pathway towards these polycyclic bioactive compounds.

Guided by the principle of parallel evolution, we identified analogous enzymes from the alkaloid-rich Solanaceae, characterising an OLADO from *N. tabacum*. The activity and alkaloid gene expression correlation of *Nicotiana* OLADO suggests it has a role in alkaloid biosynthesis, especially of anabasine, which is derived nonsymmetrically from lysine^18^. OLADOs may also contribute to other Solanaceae pathways, including nicotine formation in *N. benthamiana* (see **Table S3**) or the tropane alkaloids in *D. stramonium* biosynthesis, which are derived nonsymmetrically from ornithine^20^.

Again, using parallel evolution as a gene discovery tool, we identified OLADOs in *Artemisia annua*, a species not reported to produce alkaloids. However, alkaloids likely derived from basic amino acids are known in the *Artemisia* genus, including flavoalkaloids and long-chain alkyl derivatives, hinting that there are similar compounds present in *A. annua*^34^.

PLP-dependent enzymes capable of decarboxylative oxidative deamination are well known, including for ornithine, which can be converted to Δ^1^-pyrrolinium by bacterial ODCs^32,40^. However, the plant OLADOs characterised here are also capable of accepting lysine or arginine as substrates in an equivalent reaction. The product from arginine, γ-guanidinobutyraldehyde (GBAL), is a known plant natural product and potential precursor to GABA^37^, and has been found as a component in some alkaloid structures^38^.

The OLADOs are part of the PLP-dependent group III ornithine/lysine/arginine (Orn/Lys/Arg) decarboxylases (OLADs), and we identified OLADO paralogs which retain typical OLAD active site residues and decarboxylation activities. Previously, OLADs were known as prokaryotic enzymes, though here we have found representatives across major clades of green plants, including major crop species, and in multiple eukaryotic clades. Their absence from the majority of animal clades and Arabidopsis likely contributed to them being overlooked. Whilst the OLADOs contribute to alkaloid biosynthesis in a lineage specific manner, the conservation of the OLADs indicates an as yet undetermined role in plant primary metabolism.

Prior to this work, there were multiple examples of PLP-dependent decarboxylases gaining oxidative activities: for example, disruption of metabolic pathways in *Escherichia coli* led to the evolution of oxidative activity in the ODC SpeC in a laboratory timescale^39^. Another example are the plant aromatic amino acid aldehyde synthases which have emerged repeatedly from decarboxylases^31^. By comparing predicted structures of *F. suffruticosa* OLAD and OLADO we identified and validated key active site substitutions. These signatures enabled us to trace independently evolved OLADOs across multiple lineages. Furthermore, the OLAD/OLADO pairs were found on the genomes as tandem duplications, which is characteristic of lineage specific neofunctionalisation^30^. Therefore, alongside the OLAD/OLADO gene discovery, which has implications across plant primary and specialised metabolism, this work highlights the predictive power of parallel evolution, which, empowered by sequence information, structural prediction, and robust phylogenetic inference, can serve as a tool for gene discovery.

## Methods

### General

Chemicals and consumables used for molecular biology and biochemistry were obtained from Millipore, Invitrogen, or Merck. Substrates used in enzyme assays were obtained from Sigma-Aldrich. Lysine isotopologs were obtained from CKisotopes (UK distributor for Cambridge Isotope Laboratories). Sanger sequence verification was obtained from Eurofins Genomics or SourceBioscience. Primers were ordered from Integrated DNA technologies. Illumina sequencing was performed by Novogene, ONT sequencing by University of York Bioscience Technology Facility. Sequencing data was analysed using Geneious Prime. Figures were compiled using R-Studio, Microsoft Excel, ChemDraw, Geneious Prime, Chimera, iTOL^40^ and Adobe Illustrator.

### Plant samples

*Flueggea suffruticosa* plants (3-5 years old) were obtained from Brighton Plants (Sussex, UK), grown from seed purchased from Chiltern Seeds (Oxford, UK) in January 2014. These seeds originated from a plant in Poland, where it is a common garden ornamental. Prior to sampling, plants were grown in spring-like conditions (20 °C 16 h day, 15 °C 8 h night).

### RNA extraction

Samples (i.e. root, stem, old leaves >3 months old, young leaves < 3 months old) were isolated from plants, immediately flash-frozen in liquid nitrogen and ground in liquid nitrogen using a pestle and mortar. Roots were washed thoroughly in water prior to freezing. Frozen ground tissue (100 mg) was extracted using a standard CTAB protocol^44^ including a gDNA removal step (Invitrogen™ TURBO DNA-free Kit). The quality of the RNA was assessed using an Agilent 2100 BioAnalyzer, with all samples achieving RNA Integrity Number >8.4.

Illumina short read RNA-sequencing was performed by Novogene (UK) Ltd, enriching for polyA mRNA transcripts, and generating paired end 150 base sequencing reads on an Illumina NovaSeq 6000. Full length cDNA sequencing libraries were prepared using the Oxford Nanopore Technologies (ONT) cDNA PCR barcoding kit (SQK-PCB109), and pooled barcoded libraries were sequenced on ONT MinION and PromethION R9.4.1 flowcells (FLO-MIN106 and FLO-PRO004).

### Transcriptome assembly

The ONT reads were base-called using Guppy v5.0.11 and subjected to quality control using PycoQC to ensure the integrity and reliability of the sequencing data^41^. Concurrently, the Illumina reads underwent quality assessment with FastQC, were trimmed of adapters with Cutadapt^42^, and ribosomal RNA was removed using BBSplit.

The long-read sequences from all samples were pooled and assembled *de novo* to construct the transcriptome using Rattle^43^, a tool optimised for long-read assembly. This initial assembly was then polished with Medaka, employing the ONT reads to correct sequencing errors. To enhance the accuracy of the assembled transcriptome further, three rounds of polishing were conducted using Pilon with the combined Illumina short reads. Post-assembly, the transcripts were filtered through Evigene using the tr2aacds.pl Perl script to remove redundancy and low-quality sequences.

For annotation, the assembled transcripts were first processed with Transdecoder to identify open reading frames (ORFs). Annotation was then performed using BLASTP and BLASTX against the UniProt Swiss-Prot database to identify known protein functions. Protein domains were delineated using InterProScan^44^.

### Transcriptome quantification

For the quantification of RNA-seq reads, we employed Kallisto, a tool designed for the fast and accurate estimation of transcript abundances through pseudoalignment^45^. Initially, a Kallisto index was generated using our assembled transcriptome as a reference. Reads were then quantified against this index, employing Kallisto’s quant command with parameters tailored to our data, including 100 bootstrap samples for uncertainty estimation. The resulting output provided both estimated counts and transcripts per million (TPM) values for each transcript, facilitating downstream analyses such as differential expression.

### Transient expression in *Nicotiana benthamiana*

*Nicotiana benthamiana* plants were grown in F2+S and Calypso insecticide-treated soil in a growth room before and after infiltration (25°C on a 16/8 h day/night cycle). Plants were infiltrated with LBA4404 *Agrobacterium tumefaciens* at 5 weeks old. For preparation of LBA4404 *A. tumefaciens*, chemically competent cells (50 µL) were transformed with plasmid (200 ng) using a heat shock protocol: after mixing cells and plasmids, cells were flash frozen in liquid nitrogen, then defrosted (5 min, 37 °C), and then incubated on ice (30 min). LB media (250 µL) was added, cells were incubated (120 rpm, 2h, 28 °C), and then plated on LB agar (50 µg/mL streptomycin, 50 µg/mL kanamycin). Colonies from plates (after 2–3 days growing at 28 °C) were inoculated into LB media and incubated (50 µg/mL streptomycin, 50 µg/mL kanamycin; 28 °C, 220 rpm, 24 hours). Cells were pelleted (20 min, 4000 xg, RT) and resuspended in infiltration media (10 mM MES, 10 mM MgCl_2_, 100 µM acetosyringone, OD600 = 0.5) and incubated (RT, 50 rpm, 2 h). Cultures were infiltrated onto the abaxial side of leaves using a 1 mL syringe. One leaf per plant and three plant replicates per plasmid construct were used. For controls we used GFP vector strains (infiltration positive control) and empty vector strain (metabolite negative control). All plants were grown in 16/8 h light/dark, 25 °C, for 5 days before harvest for analysis.

Leaf discs (8 mm diameter) were excised and placed into wells (24 well plate) with buffer (400 µL, 50 mM HEPES, pH 7.5), epidermis facing upwards. Discs were gently pressed to sink to the bottom of the well, and then substrate was added (50 µL, 10 mM lysine). The wells were sealed and placed in a growth chamber for one light cycle (16 h light, 8 h dark, 25 °C). For metabolite extraction, discs were removed from the wells, dried on tissue paper, placed in 2 mL tubes and the tubes frozen in liquid nitrogen. For metabolite isolation, three tungsten carbide beads were added to each tube, and samples were homogenised to a fine powder using a TissueLyser (2x 40 s, 25 Hz) whilst frozen. Solvent was added (200 µL, 80:20 MeOH:H_2_O), tubes were vortexed (1 h, RT), and samples centrifuged (10,000 xg, 5 min). Extracts were diluted with MeOH (600 µL), filtered through 0.45 µm PTFE filters and transferred to LC-MS glass vials for subsequent analysis.

### Protein expression and purification

For protein expression, *E. coli* Rosetta™(DE3) or SoluBL21 (DE3) (Genlantis) were used. Single colonies transformed with plasmids containing genes of interest were used to inoculate overnight cultures with antibiotic selection (2xYT media, 37 °C, 200 rpm). These were diluted with 2xYT with antibiotics (OD600 = 0.1) and grown to OD600 = 0.7 for Rosetta cells or OD600 = 0.4-0.6 for SoluBL21 (37 °C, 200 rpm). Then the cells were induced with IPTG (1 mM) and incubated overnight (∼14 h, 200 rpm, 18 °C). Cells were harvested by centrifugation (3200 xg, 10 min, 4 °C), the supernatant removed, and the pellet resuspend in lysis buffer (10% v/v compared to original culture; 50 mM Tris-HCL, 50 mM glycine, 5% v/v glycerol, 0.5 M NaCl, 20 mM imidazole, pH 8; per 50 mL, 1x EDTA-free protease inhibitor and 10 mg lysozyme) and incubated (30 min, 4 °C). The cells were lysed through a cell disruptor (26 kPsi) and clarified by centrifugation (20 min, 35,000 xg, 4 °C). Purification was achieved through nickel affinity purification using an ÄKTA start FPLC. Clarified lysate was loaded onto a Ni-NTA column (5 mL) and then the column washed with binding buffer (50 mM Tris-HCL, 50 mM glycine, 5% v/v glycerol, 0.5 M NaCl, 40 mM imidazole, pH 8) and the protein eluted (50 mM Tris-HCL, 50 mM glycine, 5% v/v glycerol, 0.5 M NaCl, 500 mM imidazole, pH 8). A protein purification outcome was inspected by SDS-PAGE, and if a pure was protein obtained, samples were buffer exchanged into 50mM HEPES, pH 7.5 using a PD10 column, using a gravity method as per manufacturer instructions. The protein was concentrated by centrifugation (Amicon^®^ Ultra-2 mL Centrifugal Filters, 30 kDa MWCO) prior to addition of glycerol (10% v/v), flash freezing and storage at –70 °C.

### Cloning

Initially, genes of interest were identified using the transcriptome assembly and cloned from the plant source. RNA isolated from *Flueggea suffruticosa* (see above) was reverse transcribed to cDNA using Invitrogen™ SuperScript IV kit as per the manufacturer’s protocol. The cDNA was immediately used for PCR amplification. Primers were designed that contained an overhang for cloning into vectors (supplementary for primers). PCR was performed using SuperFi™ polymerase enzyme and amplified genes were isolated using a gel extraction kit. For LaLDC and truncated PS/PSP sequences in *E. coli*, codon-optimised sequences in pET-28a(+) were obtained using gene synthesis (Twist).

Purified pHREAC vector plasmid^46^ was digested with BsaI (50 µL reactions; 1 µg DNA, 10 U of BsaI in CutSmart™ Buffer, 37 °C, 1 h) and processed using a PCR clean-up kit (Promega Wizard™). Amplified genes with pHREAC overhangs, deriving either from the cDNA or a plasmid (see above), were inserted into linear pHREAC using In-Fusion HD cloning (with insert:plasmid 2:1). A colony PCR using pHREAC primers (see supp table) was carried out to confirm plasmid transformation and successful cloning. After overnight incubation of transformed *E. coli*, plasmids were miniprepped and sequence verified before transformation into LBA4404 *Agrobacterium tumefaciens*.

### Mutagenesis

Forward and reverse site directed mutagenesis primers were designed to overlap by 8-15 bases at the 5’ end, with the mutation present in the overlapping region^47^. The templates were the codon optimised genes on pET-28a(+). Primers were synthesised by Integrated DNA Technologies UK LTD. Plasmids were propagated in DH5α *E. coli* cells and extracted using the Zymopure™ plasmid miniprep kit. PCR was performed using SuperFi™ polymerase enzyme. Each PCR reaction (50 μL) mix contained template DNA (0.2 ng/μL), DMSO (1.5% v/v), Platinum™ SuperFi II PCR Master Mix (50% v/v), and forward and reverse primers (0.5 μM). PCR cycles were initiated at 98°C for 1.5 min; followed by 30 amplification cycles: 98 °C for 10 sec, 60 °C for 10 sec and 72 °C for 3.5 mins; then an elongation step at 72 °C for 5 min. DpnI (1 μL) restriction enzyme was added directly to the PCR mix and left to incubate overnight at room temperature. PCR mix was used to transform into DH5α E. coli cells to propagate plasmid and extracted using the Zymopure ™ plasmid miniprep kit. Plasmids were sequence verified before transformation into SoluBL21 (DE3) (Genlantis) for protein expression and purification.

### *In vitro* enzyme Assays

Reaction components were mixed (100 nM, 10 ng/µL or 20 ng/µL purified protein, 0.1 mM PLP, 50 mM NaCl, 2 mM DTT, 50 mM HEPES pH 7.5, 5 or 10 mM substrate, 100 μL final volume). Substrate was added last and the reactions incubated (16 h, 30 °C) prior to quenching with MeCN (100 or 400 μL) and LC-MS analysis. For protein controls, proteins were pre-incubated (5 min, 99 °C) prior to substrate addition. Reactions were performed in triplicate, with similar outcomes. One representative chromatogram is presented. For wild-type assays, the substrate was at 10 mM whereas the FsOLAD/OLADO substitution assays used 5 mM substrate.

### LC-MS analysis

Enzyme activity analysis was performed using hydrophilic interaction liquid chromatography (HILIC). Chromatography was performed using a HPLC instrument Dionex UltiMate 3000 RSLC nano system or a Thermo Fisher Vanquish UHPLC. The column was an XBridge^®^ BEH Amide column (Waters, 5 µm, 4.6mm x 100 mm). Phase A [H2O 0.125% v/v formic acid, 10 mM ammonium formate] and phase B [95:5 MeCN:Phase A] were used as the mobile phase components. For the Dionex, chromatography was performed with 0.6 mL/min flow rate and a gradient: 0-4 min, 100% B; 4–24 min, 100–60% B; 24–25 min, 60–100% B; 25–35 min, 100% B. For the Thermo Fisher, chromatography was performed at 0.6 mL/min flow rate and 30 °C with gradient: 0-2 min, 100% B; 2-10 min, 100-60% B; 10-11 min, 60-100% B; 11-12 min, 100% B. Mass spectrometry was performed with an MS Bruker HCT ultra-ETD II (50-500 m/z collected). Standards were dissolved in either phase A or B depending on solubility. Data was processed in Bruker Compass DataAnalysis Version 4.4 (Dionex) or Freestyle (Thermo).

### H_2_O_2_ Assays

Peroxide formation was detected using phenol and 4-aminoantipyrine (4-AAP) to produce colorimetric readout at 505 nm. Reactions contained: 0.5 mg/mL purified enzyme, PBS pH 8.0, 20 mM lysine, 1 mM DTT, 4 mM phenol, 6 mM 4-AAP, 100 U peroxidase and PLP (0.0125-0.2 mM). Negative control assays were performed without enzyme, and positive controls by the addition of H_2_O_2_. Spectrophotometric measurements were taken using a CLARIOstar^®^ Plus UV-Visible spectrophotometer (BMG Labtech) at 505 nm in continuous plate mode. Runs were performed in triplicate.

### 13C-NMR Assays

Enzyme assays performed as previously described but with ε^15^N, ε^13^C-lysine or ^13^C_6_-lysine. After incubation (16 h, 30 °C), samples were pooled, diluted in 10:90 D_2_O:H_2_O and place in an NMR tube (S400). The spectrum was measured at 700 MHz using a Bruker NEO spectrometer equipped with a cryoprobe prodigy TCI. 13C spectra used 14,000 scans with a 2s relaxation delay. Data analysis was carried out using MestRe Nova V14.

### Synthesis of Δ^1^-piperideine

To a solution of *N*-chlorosuccinimide (8.34 g, 62.5 mmol) in dry diethyl ether (60 mL) under argon was added piperidine (5.90 mL, 58.7 mmol) dropwise and the resulting mixture was stirred for 2 hours at room temperature. The reaction mixture was then filtered and the filtrate was washed with water (2 x 35 mL), dried (Na_2_SO_4_), filtered and concentrated *in vacuo* to give an oily residue. A solution of NaOMe in MeOH, freshly prepared by adding sodium metal portionwise (1.53 g, 66.7 mmol) to MeOH (35 mL), was added to the oily residue at room temperature. The reaction mixture was stirred at reflux for 45 mins. After cooling to room temperature, water (50 mL) was added and the mixture was extracted with Et_2_O (3 × 50 mL). The combined organic layers were dried (MgSO_4_), filtered and concentrated *in vacuo* to give Δ^1^-piperideine (2.72 g, 56%). NMR data obtained were consistent with the literature. NMR data obtained matched those previously described^48^.

### Synthesis of Δ^1^-pyrrolinium

To form Δ^1^-pyrrolinium *in situ* for LC-MS analysis, 4-aminobutyraldehyde diethyl acetal (SigmaAldrich) was dissolved in 0.1 M HCl and incubated at 30 °C for 30 mins prior to analysis.

### Synthesis of 4-guanidinobutyraldehyde diethylacetal hemisulfate

To a solution of *S*-methylisothiourea hemisulfate (0.278 g, 2.00 mmol) in water (4 mL) and ethanol (4 mL) was added 4-aminobutyraldehyde diethyl acetal (0.345 mL, 2.00 mmol) and the reaction mixture was stirred at room temperature for 18 hours. The reaction mixture was then concentrated *in vacuo*. Purification by trituration with diethyl ether gave 4-guanidinobutyraldehyde diethylacetal hemisulfate (0.423 g, 84%) as a white solid. Mp 95–97 °C; *ν*_max_/cm^-1^ (neat) 3139, 2973, 2881, 1689, 1634, 1372, 1062, 596; *δ*_H_ (400 MHz, DMSO-*d*_6_) 1.18 (6H, t, *J* 7.1 Hz), 1.59–1.84 (4H, m), 3.12–3.23 (2H, m), 3.47–3.57 (2H, m), 3.61–3.72 (2H, m), 4.53 (1H, t, *J* 5.2 Hz); *δ*_C_ (101 MHz, DMSO-*d*_6_) 15.7 (2 × CH_3_), 25.3 (CH_2_), 31.9 (CH_2_), 42.1 (CH_2_), 62.8 (2 × CH_2_), 104.2 (CH), 158.7 (C); *m/z* (ESI) 204.1708 (MH+. C_9_H_22_N_3_O_2_ requires 204.1707). To form 4-guanidinobutyraldehyde *in situ* for LC-MS analysis, the diethylacetal hemisulfate was dissolved in 0.1 M HCl and incubated at 30 °C for 30 mins prior to analysis.

### Protein structural analysis

Structural models of enzymes were constructed using AlphaFold2 and AlphaFold2-multimer on ColabFold^49^. PLP cofactors were added with AlphaFill^50^. The best scoring model was selected for display. For dimers of PS, the unstructured N-terminal sequence was removed prior to calculation. The structural models, along with X-ray structures were depicted using Chimera (UCSF). For EcLDCI (PDB: 3N75) selected subunits of the homo 10-mer were depicted to enable visual comparison.

### Phylogenetic analysis

For phylogenetic analysis of the group III decarboxylase proteins, sequences of proteins containing the Orn/Lys/Arg decarboxylase, major domain were obtained from InterPro (IPR000310). Of bacterial sequences, only those which were Swiss-Prot “Reviewed” were kept. All eukaryotic sequences were selected, along with Fs1864/Fs3498 from *F. suffruticosa*, and the group II decarboxylase outgroup. Then duplicate and obsolete sequences were removed, as were sequences with <400 residues and those without Pfam PF01276 OKR_DC_1 annotation. After alignment with MAFFT^51^, sequences were further curated by removing those missing the PLP binding lysine and the conserved glycine (570 on PS), selecting a single sequence from those with >95% sequence identity with others, and removing sequences with large alignment gaps with respect to PS residues 98-580. A maximum likelihood tree was then inferred using W-IQ-TREE^52^ with ModelFinder^53^, ultrafast bootstraps (UFBoot2, X1000)^54^, and SH-aLRT supports (X1000)^55^. For the rosid phylogeny, selected sequences were obtained, realigned with MAFFT, then columns with >95% gaps were removed prior to tree inference with W-IQ-TREE. Trees were depicted using iTOL^40^.

For the focussed asterid phylogeny, we accessed published genomes to obtain homologs from: *Nicotiana tabacum*^56^, *Artemisia annua*^57^, *Atropa belladonna*^58^, *Datura stramonium*^58^, *Solanum lycopersicum* (ITAG4.1)^59^, *Capsicum annuum* (UCD10X, Ref), *N. benthamiana* (Niben1.0.1)^59^ and *N. attenuata* (NIATTr2)^60^. The nucleotide sequences from *Nicotiana tabacum* homologs identified from the InterPro domain were used as queries for blastn or Megablast with default settings, with the genome extracted coding sequences (CDS) as the databases. The top hit genes according to blastn or MegaBlast, as judged by Bit-Score were obtained, filtered for length (>1000 nucleotides), translated to peptides and then used in the alignment. The alignments were performed with MAFFT and the tree inferred with W-IQ-TREE. For *N. tabacum* and *A. annua* the genomic location of the genes of interest were viewed using the genome fasta and gff file in Geneious Prime 2024.

### *N. benthamiana* gene expression

For *N. benthamiana* the gene expression was assessed using the Plant Gene Expression Omnibus ^61^. For gene expression boxplots the TPM values across all 476 samples in the omnibus were downloaded for each OLAD/OLADO paralog and plotted in R-Studio, grouping by tissue. For expression correlations, we used the inbuilt co-expression calculation (PCC value) with the exception of the ODC genes, for which we calculated PCCs manually. *N. benthamiana* alkaloid biosynthesis gene orthologs were annotated using *N. tabacum* derived query sequences and obtaining top hits from tblastn on the extracted coding sequences (CDS) from the *N. benthamiana* genome. UniProt accession values for queries: PMT1_TOBAC (PMT, putrescine methyltransferase), AO2A_TOBAC (AO, aspartate oxidase), IFRH_TOBAC (A622), ODC1B_TOBAC (ODC, ornithine decarboxylase [group IV]), QPT2B_TOBAC (QPT, quinolinate phophoribosyltransferase), QS1_TOBAC (QS, quinolinate synthase), F1T160_TOBAC (BBL, berberine bridge enzyme), and MPO1_TOBAC (MPO, N-methylputrescine oxidase).

## Acknowledgements

We thank Matthew Davy for running samples on the Jeol-700 NMR. We also thank Dr Ed Bergstom for his training and guidance with the LC-MS instrumentation, and the Centre for Excellence in Mass Spectrometry for equipment. The plasmid pHREAC was a gift from George Lomonossoff (Addgene plasmid #134908). We are grateful for computational support from the University of York High-Performance Computing service, Viking, and the Research Computing team. B.R.L. acknowledges support from UKRI (MR/S01862X/1) and BBSRC (BB/Y003586/1).

## Author Contributions

C.X.W. carried out the nucleic acid extraction, transcriptome analysis, cloning, enzyme purification/characterisation and chemical synthesis. Z.J., U.S.B and I.A. performed enzyme purification and characterisation. I. A. performed mutagenesis. L.J.N.W. performed chemical synthesis. O.S.D. performed the peroxidase assays. S.J. prepared sequencing libraries and ran ONT sequencing. K.N. assembled the transcriptome. W.P.U. assisted in the chemical synthesis and interpretation of NMR data. G.G. assisted in the interpretation of data. B.R.L. designed and supervised the project and conducted the phylogenetic and structural analyses. C.X.W. and B.R.L. designed the experimental procedures, analysed data, prepared figures, and wrote the manuscript.

## Data availability

Raw read sequences for all generated data in this study are available in the National Center for Biotechnology Information Sequence Read Archive under BioProject ID PRJNA1106657. Sequences of coding sequences of genes are available on GenBank: gene 1864 (FsOLADO) (PP748646), gene 3498 (FsOLAD) (PP748647), gene 4984 (PP748648) and gene 4074 (PP748649). All other data is available upon reasonable request to the corresponding author.

