## Supplementary Information for "Parallel evolution of plant alkaloid biosynthesis from bacterial-like decarboxylases"

Supplementary Figures S1-S12 Page 2

Supplementary Tables S1-S5 Page 13

Supplementary References Page 19

### Supplementary Figures

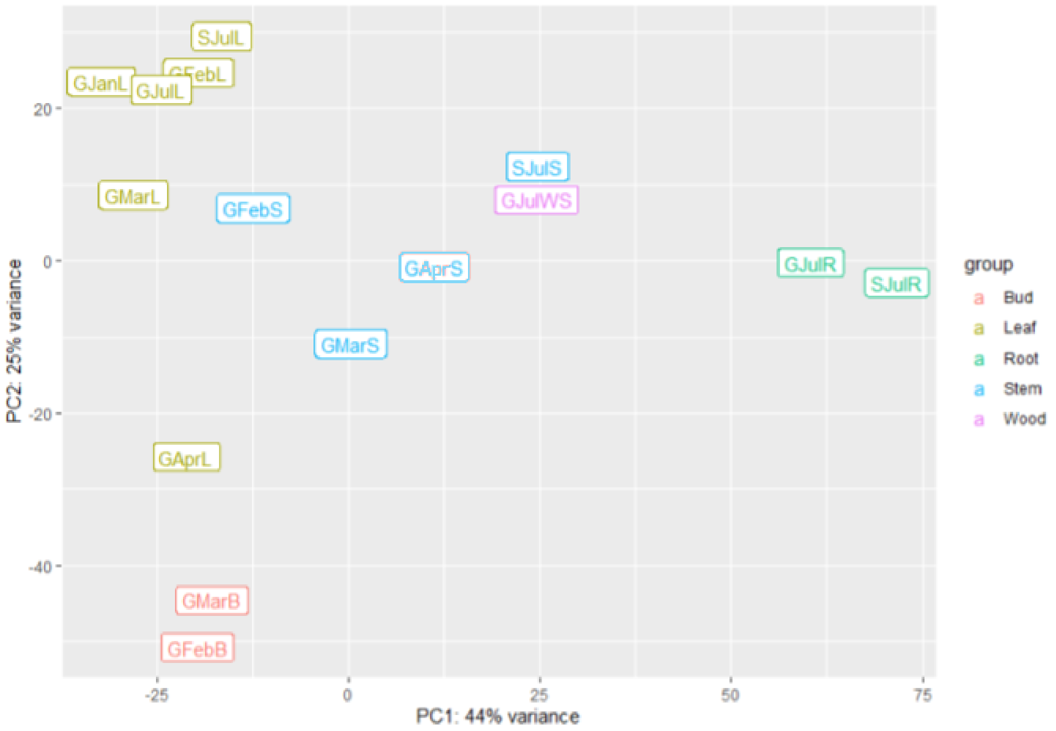


**Figure S1. Principal component analysis of RNAseq samples.** Principal component analysis (PCA) plot of all 15 samples sequenced. The plot positions samples based on their transcriptional profiles along the first principal component (PC1) on the x-axis, which captures the greatest variance, and the second principal component (PC2) on the y-axis, representing the next greatest variance. Codes for samples in **Table S1**.

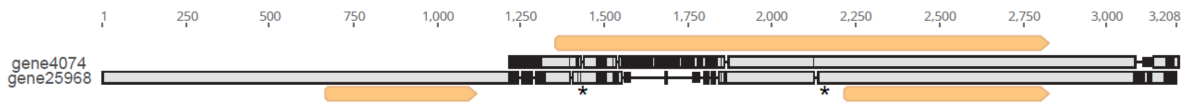


**Figure S2. Alignment of two lysine decarboxylase candidates.** Nucleotide alignment of genes 4047 (top) and 25968 (bottom). Grey boxed regions are identical, black regions with corresponding lines show gaps. Orange annotations are open reading frames (ATG start, minimum 400 nucleotides). Stars on 25968 show stop codons in frame with long ORF from 4074. The long ORF from gene 4074 was selected as the sequence to be screened.

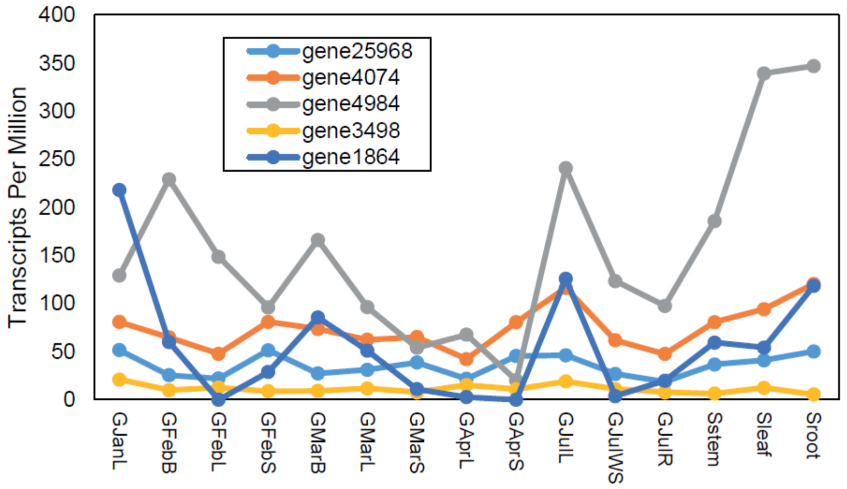


**Figure S3. Expression levels of LDC candidate genes.** RNAseq determined expression levels across 15 samples. Codes for samples in **Table S1**.

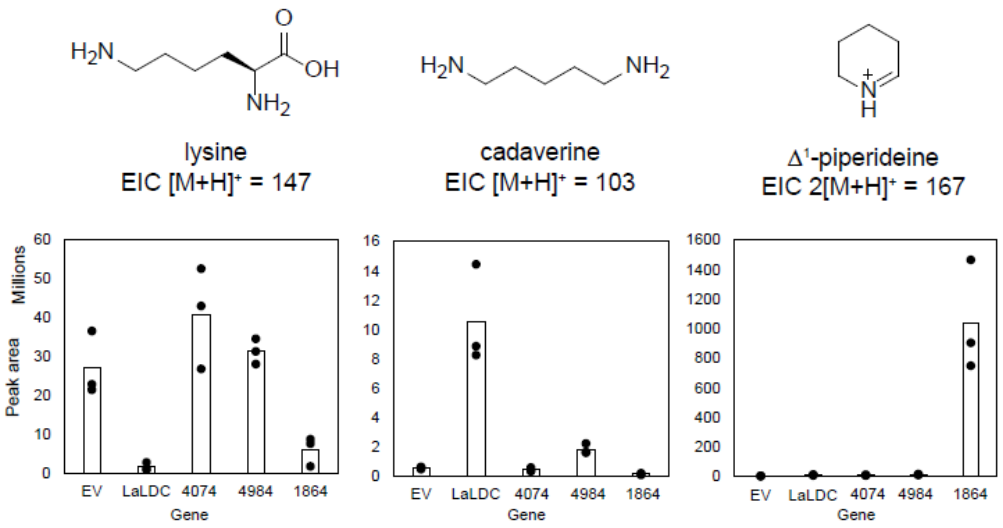


**Figure S4. Screening of lysine decarboxylases.** LC-MS analysis of leaf disc assays (see **Fig. 2B**). Bar graphs show corresponding EIC peak areas of lysine, cadaverine and Δ[1]-piperideine; bars show mean of triplicate reactions, points show from each reaction. All areas were blank corrected to the average area of the non-infiltrated plants. *Lupinus angustifolius* LDC (LaLDC) is positive control, empty vector (EV) is negative control.

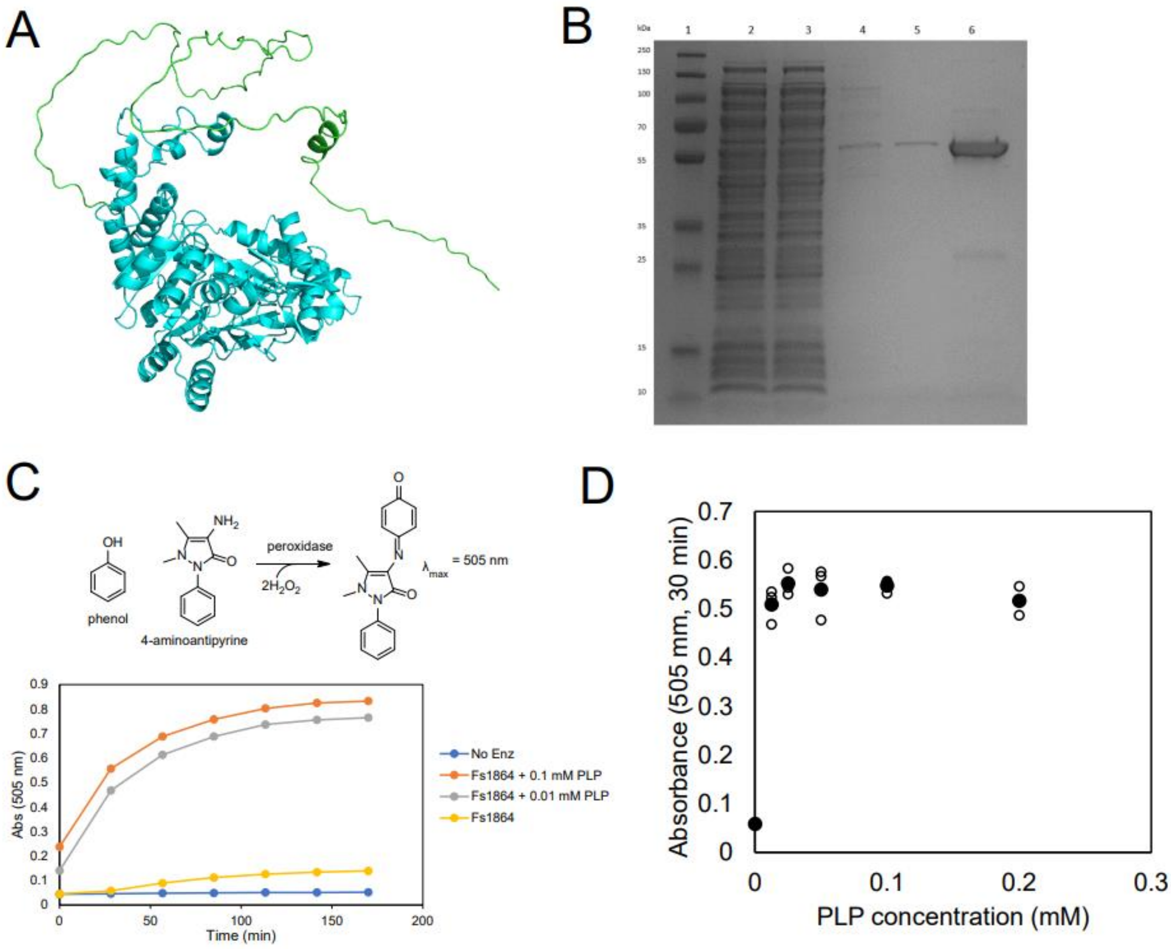


**Figure S5.** ***In vitro*** **assessment of Fs1864 activity. A.** AlphaFold structural model of Fs1864 monomer with long N-terminus highlighted in bright green and globular fold in aqua. The green region (74 amino acids) was removed for expression in *E. coli.* **B.** SDS-PAGE showing purified Δ74N-PS. Lanes: 1, ladder; 2, clarified lysate; 3, flow-through; 4, wash 20 mM imidazole; 5, wash 40 mM imidazole; 6, 500 mM elution. **C.** Coupled enzyme assays showing $H_2O_2$ production in Fs1864. **D.** Dependence of $H_2O_2$ production on PLP concentration, determined by peroxidase coupled assay. Solid circles are means of triplicate measurements. Individual measurements are in hollow circles.

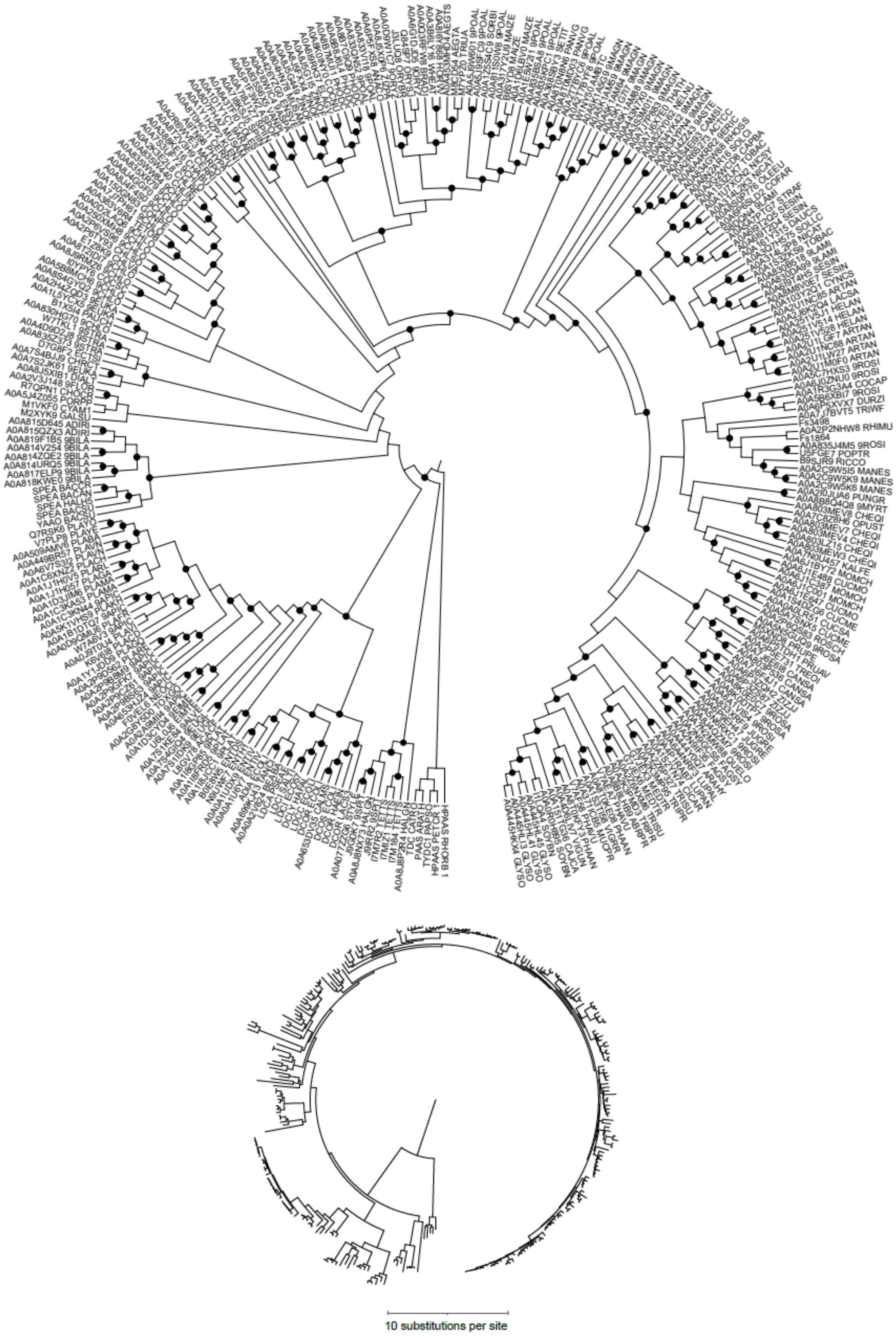


**Figure S6. Maximum likelihood phylogeny of group III decarboxylases.** Replication of tree in **Fig. 5B** but with Uniprot accession labels (top) and branch lengths show (bottom). In the top tree, black dots on nodes show highly supported branches (SH-aLRT ≥80% and UFboot ≥95%).

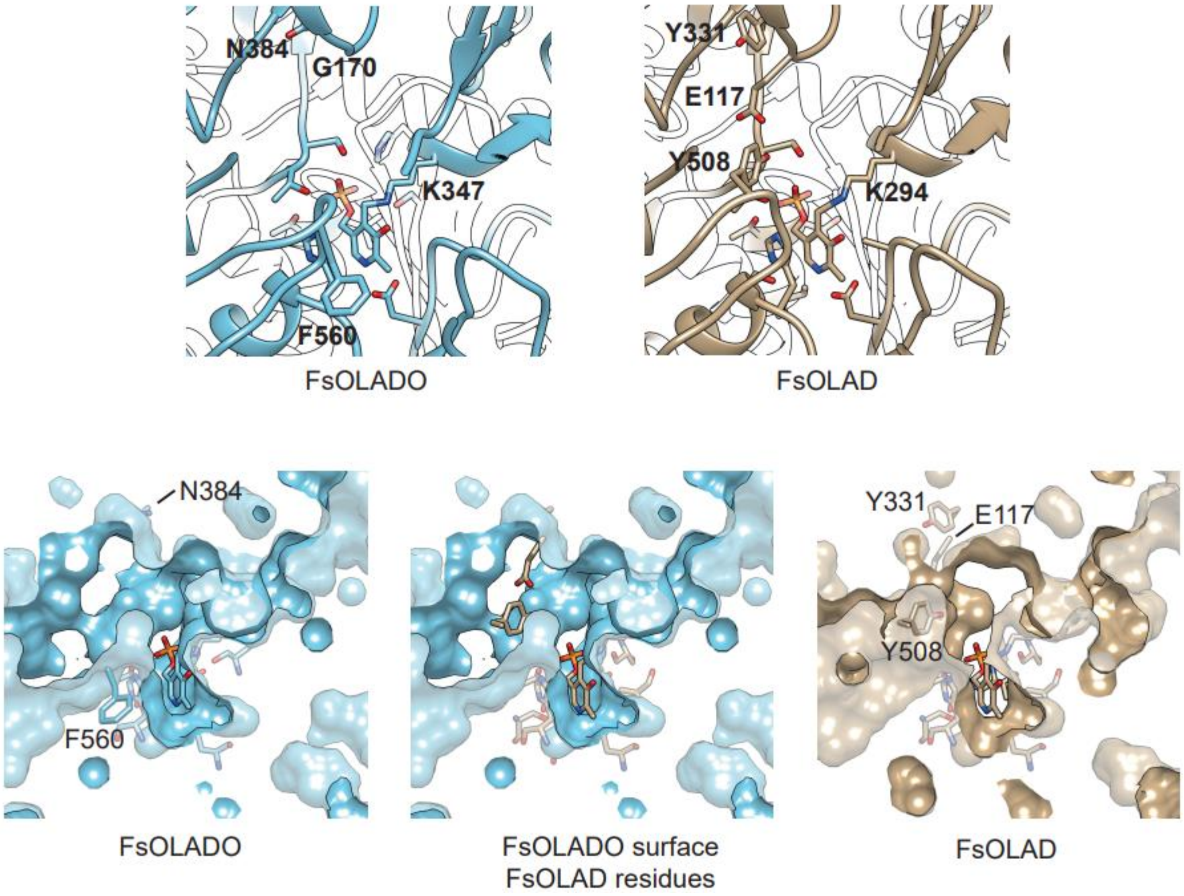


**Figure S7. Structural models of FsOLAD and FsOLADO.** Predicted dimer structures of FsOLADO (blue) and FsOLAD (beige), predicted with AlphaFold and PLP added with AlphaFill. Top depiction showing backbone as ribbon with depth cueing (transition from colour to clear). Selected residue side chains depicted and labelled. Bottom panels show active site region with selected residues depicted and shown with a clipped-slab section of predicted molecular surface, highlighting size of active site. Centre shows OLADO surface on OLAD residues after structural alignment, highlighting the active site size implication of the OLAD/OLADO E117/G170 and Y508/F560 substitutions. Depiction performed in UCSF Chimera.

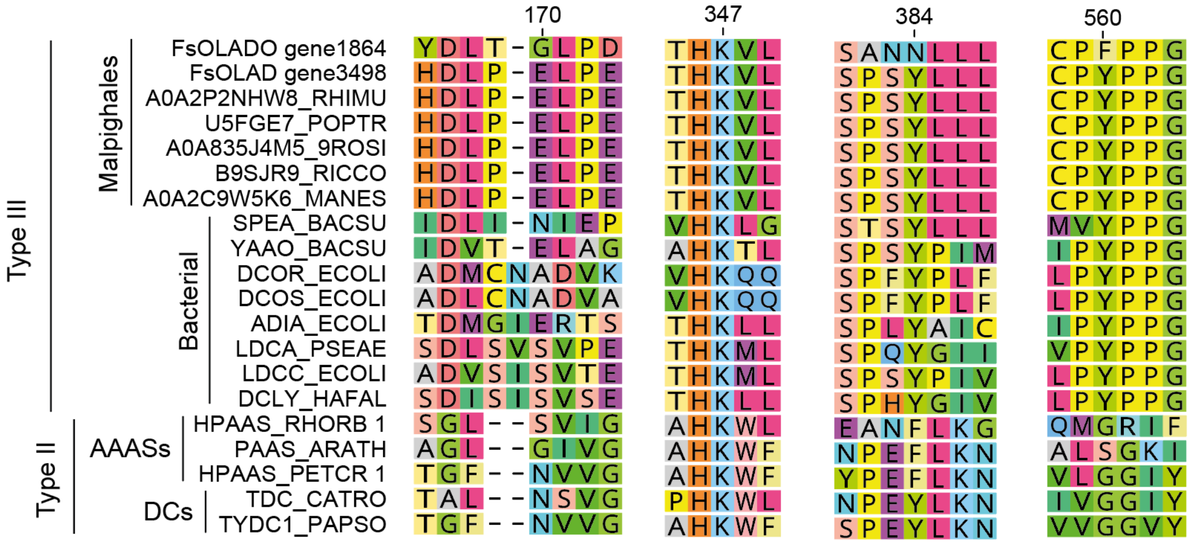


**Figure S8. Sequence alignment of PLP-fold type I decarboxylases.** Selected aligned regions and sequences showing notable residues. Residue numbering is based on FsOLADO. AAAS = amino acid aldehyde synthase, DC = decarboxylase. Types refer to PLP-dependent decarboxylase types.

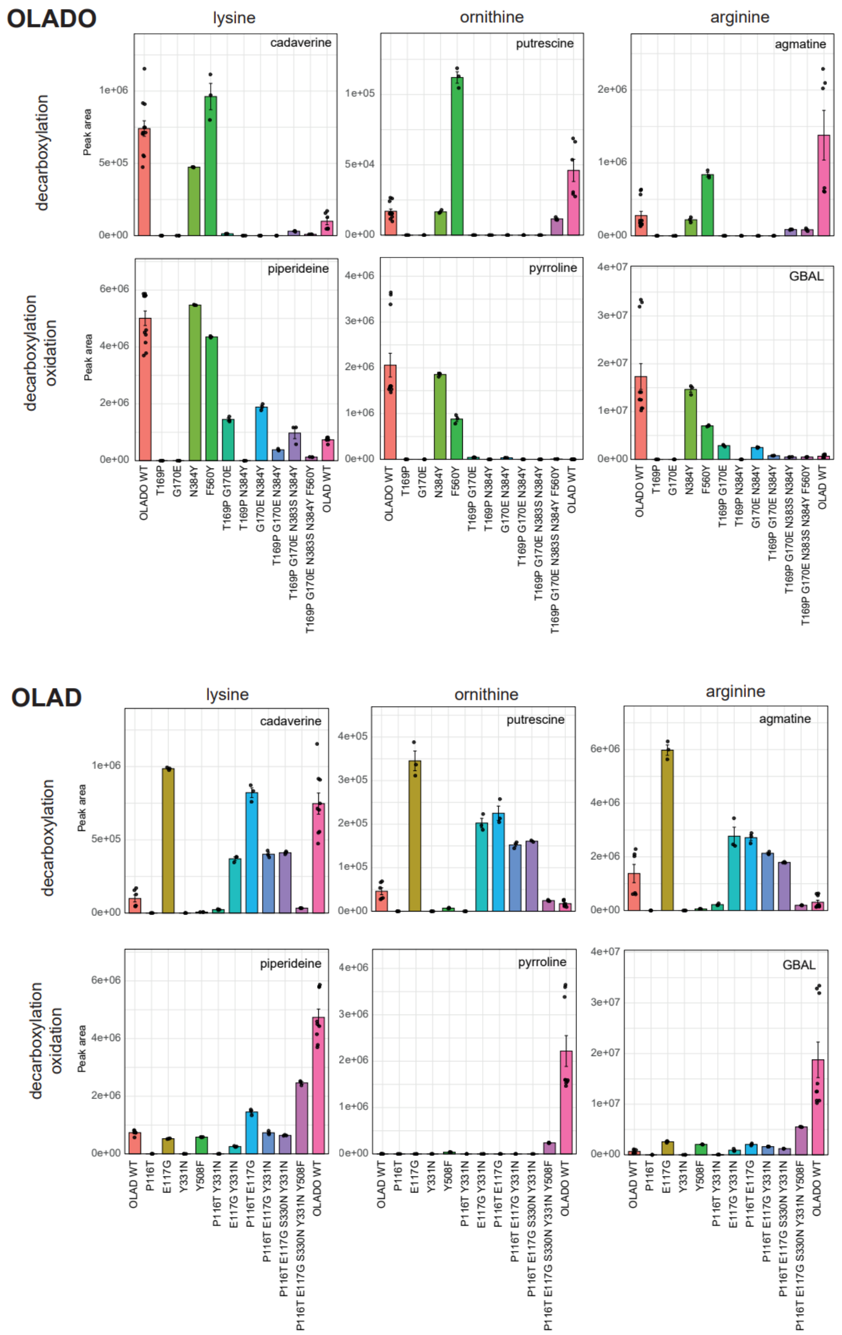


**Figure S9. OLAD/OLADO variant activities.** Activity of purified enzymes with lysine, ornithine and arginine. LC-MS analysis of *in vitro* assays, forming either the decarboxylated product (cadaverine, putrescine or agmatine) or decarboxylated and oxidised (and cyclised) product ($\Delta^1$-piperideine, $\Delta^1$-pyrrollinium and γ-guanidinobutyraldehyde [GBAL]). Bar graphs show corresponding LC-MS derived EIC peak areas; bars show mean of multiple reactions, points data from each reaction, error bars are SE. Substrate concentration 5 mM, enzyme concentration 20 ng/µL. See **Fig. 4C** for the heatmap version of this data.

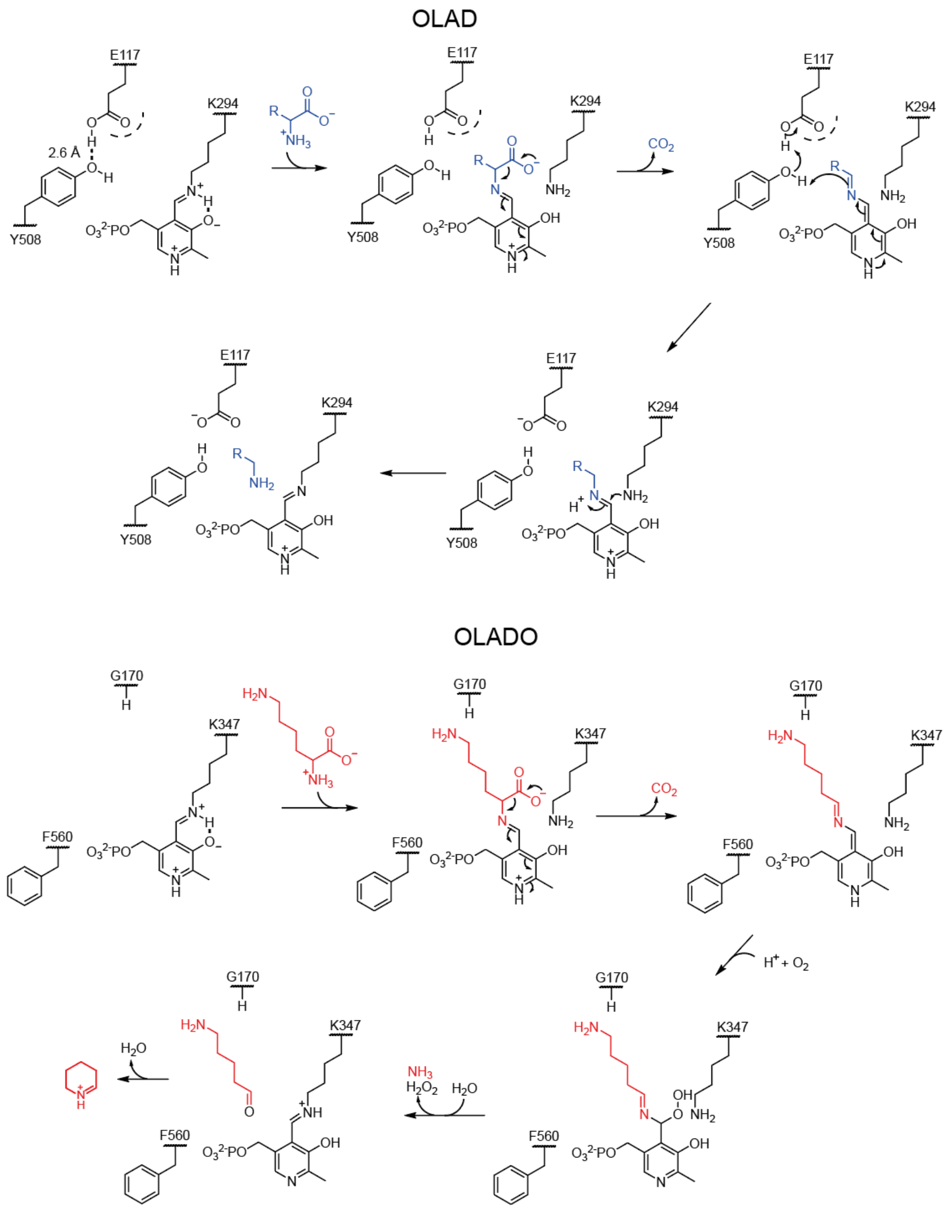


**Figure S10. Proposed OLAD/OLADO enzyme mechanism.** Proposed mechanism of OLAD decarboxylation (top, with generic amino acid) and OLADO decarboxylation-oxidation (bottom, with lysine) highlighting the possible roles of Y508/F560 and E117/G170 in determining reaction and substrate acceptance. The scheme is based on that in Hoffarth *et al* 2020[1] and Torrens-Spence *et al* 2020[2].

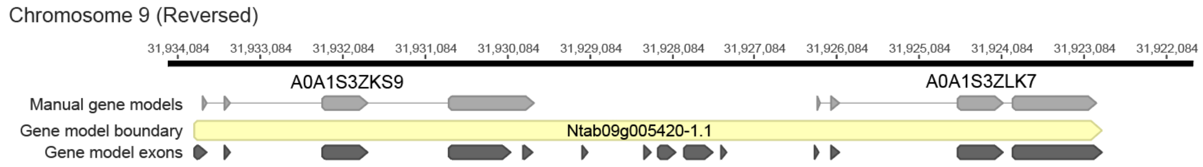


**Figure S11. *Nicotiana tabacum* OLAD/OLADO.** Details of manual correction of gene models of Ntab09g005420. The UniProt based sequences originate from Sierro *et al* 2014[3]. The gene model fusion is indicative of a misannotated tandem duplication event.

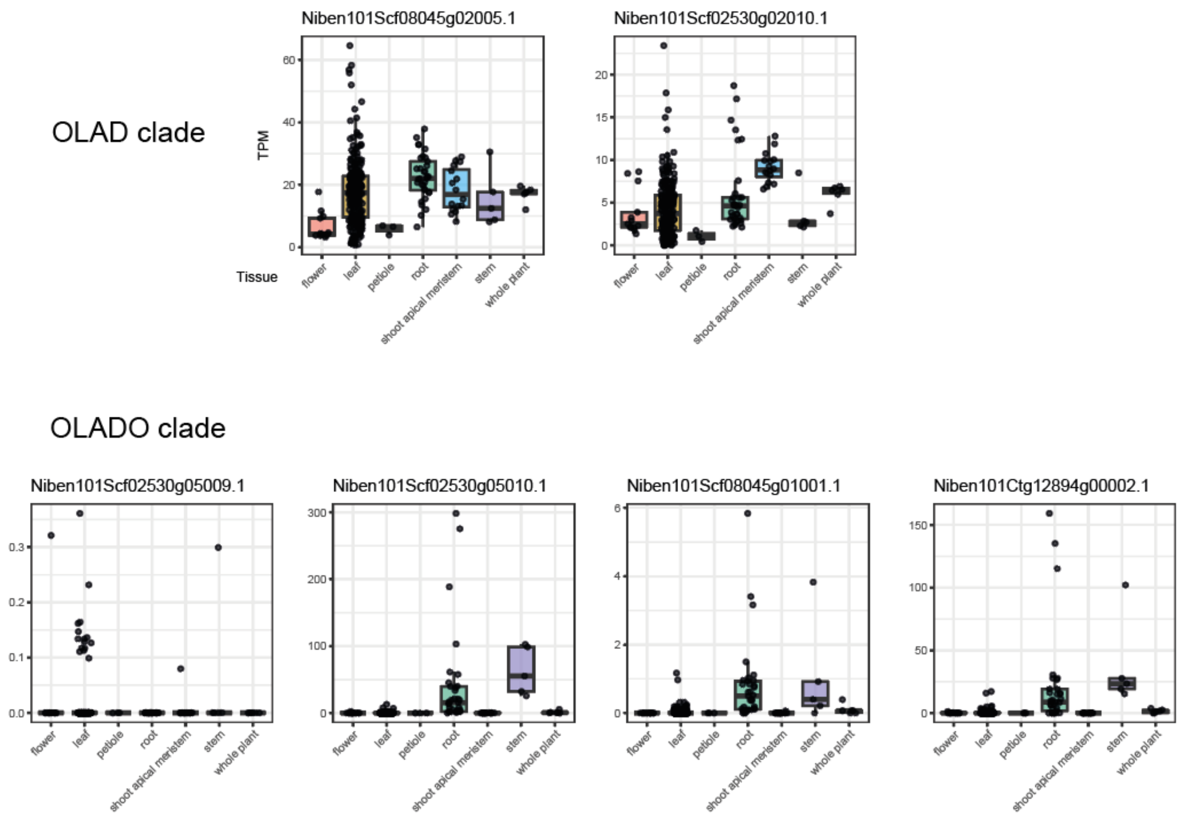


**Figure S12. *N. benthamiana* OLAD/OLADO expression.** Expression (TPM) of *N. benthamiana* (Niben1.0.1) OLAD/OLADO homolog genes as identified by top blast hit using *N. tabacum* genes as queries. Fragmentary genes' expression is not shown, with the exception of Ctg12894g00002 which was included due to high correlation with Scf02530g05010 (**Table S3**). Expression calculated across 476 samples, grouped by tissue type. Compiled from the Plant Expression Omnibus database[4].

**Supplementary Tables**

**Table S1. RNA Samples.** *Flueggea suffruticosa* RNA samples used in the study for *de novo* transcriptome and gene expression analysis. Sequencing available on NCBI (BioProject PRJNA1106657)

| Sample Number | Sample code | Plant code | Month | Tissue | Sequencing technology |
|---|---|---|---|---|---|
| 1 | GJanL | G | January | Leaf | Illumina PE150 |
| 2 | GFebB | G | February | Bud | Illumina PE150 |
| 3 | GFebL | G | February | Leaf | Illumina PE150 |
| 4 | GFebS | G | February | Stem | Illumina PE150 |
| 5 | GMarB | G | March | Bud | Illumina PE150 |
| 6 | GMarL | G | March | Leaf | Illumina PE150 |
| 7 | GMarS | G | March | Stem | Illumina PE150 |
| 8 | GAprL | G | April | Leaf | Illumina PE150 |
| 9 | GAprS | G | April | Stem | Illumina PE150 |
| 10 | GJulL | G | July | Leaf | Illumina PE150 |
| 11 | GJulWS | G | July | Woody stem | Illumina PE150 |
| 12 | GJulR | G | July | Root | Illumina PE150 |
| 13 | Sstem | S | July | Stem | Illumina PE150 |
| 14 | Sleaf | S | July | Leaf | Illumina PE150 |
| 15 | Sroot | S | July | Root | Illumina PE150 |
| | Pooled samples | | | | ONT MinION |
| | Pooled samples | | | | ONT PromethION |

**Table S2. Cloning primers.** Used to cloned candidates in pHREAC vector. Italics represent homolog overhangs.

| Gene Candidate | Direction | Sequence |
|---|---|---|
| 4074 | Forward | *TATATTAAACGTCTCTAAAAATG*GCGGCGACAACGTCTC |
| 4074 | Reverse | *AATTTAATGAAACCAGAGCG*TCACAGGCCTTCGAAGAATCG |
| 4984 | Forward | *TATATTAAACGTCTCTAAAAATG*CCGGCCCTGGCGTGTTG |
| 4984 | Reverse | *AATTTAATGAAACCAGAGCG*TCAAGCACAGCAATAGGACC |
| 1864 | Forward | *TATATTAAACGTCTCTAAAAATG*GCTAACAATCAGACTCTCAG |
| 1864 | Reverse | *AATTTAATGAAACCAGAGCG*CTACACATCACATATGACTATAG |

**Table S3. Amino acid sequences.** Used for gene synthesis. pET28 vector, codon optimised. Met sometimes present

| Species | Protein name | Source | Amino acid insert |
|---|---|---|---|
| *Flueggea suffruticosa* | Fs1864 (OLADO) | *De novo* transcriptome (this work) | TTTPFLSALKAAAEKNAASFHFPGHNRGHAAPPSLIDLIGLRPFMYDLTGLPDIDALSCPKGPILEAQQ<br>EAAKLFGASETWFLVGGTTCGVLAAIMGTCSPGDHIILTRNCHVSAISAMVLCGAVPKYMIPEYNSNW<br>DVAGGITLSQVKEAIEELEMEGQKPAAIFITSPTFQGICSNIGEISRLCHSYGIPLIVDEAHGAHLGFHPD<br>MPRSALSQGADLVVQSTHKVLPALSQSSMLHISGNFVSRDRISRCLQILQSTSANNLLLASLDAARAQ<br>LAENPIDIFEEPLDIAREARSLIKNIPHIGLLGLETFPKFEEMDPMRLTVGFWQLGLSGYEACEILERHQ<br>GVVPEMIGTKSITLPFNLGTRRDHVRRLAAGLEELSARSFQVAGMDKSVVTEGHEVFADVTVHLNPR<br>DAFFAKKRTVSIRESLGKVCGELICPFPPGIPVMIPGEIITEKVLNYILDVKRKGIVVSGASDPLLTTIVIC<br>DV |
| *Flueggea suffruticosa* | Fs3498 (OLAD) | *De novo* transcriptome (this work) | GATPLVSALKAAAEKDAASFHFPGHNRGRAAPSSLTQLIGVRPFLHDLPELPELDNLFSPEGPILEAQ<br>KKAAKLFGAKETWFLVGGTTCGIQAAIMATCSPGDYIVLPRNSHMSAISGIVLSGAIPKYIIPQYDIDWDI<br>AGGVSPSQVEKAITELQFDGRKPAAIFVTSPTYHGICSNLSEISQLCHSYGIPLIVDEAHGAHFGFHPQ<br>MPHSALQQGADLVVQSTHKVLCSLTQSSMLHMVGDIIDRERIFWCLQTLQSTSPSYLLLASLDAARSQ<br>ISENPDSVFNQAMKLSSEAKACIKNIPGITVLDFSSFDKFPIIDPLRLTIGFWQLGLSGYEADDILEREYG<br>IVSELVGSKSITFAMNLGTCREHVERLETGLKRLSACSFQNEQLENRTKKGYDYEPFSDFVMSMNPR<br>DAFFSRKRRVSIEESIGKISGELVCPYPPGIPILIPGETITQNALDYLLDCRNKGAVVTGASDPLLSSLLIC<br>DV |
| *Nicotiana tabacum* | NtOLAD A0A1S3ZLK7 | UniProt / Sierro *et al* 2014 genome[3] | ELPPLVSALKASAEENAASFHFPGHNRGKAAPSSLTQLIGSKPFLHDLPELPELDNLFSPEGPILEAQK<br>QAARLFGASETWFLVGGSTCGIQAAIMATCSPGDTLILPRNSHISAISSMVLSGTLPKYIVPEYDFNWD<br>VAGGITPSQVHKAIKELEMEGRKAAAVLVVSPTYHGICSNLDEICEICHSHSIPVIVDEAHGAHLGFHLK<br>LPSSSLSQGADLSVQSTHKVLCALTQSSMLHMQGNLVDRERISRSLQMLQSSSPNYLLLASLDAARA<br>QISENREAIFYQTIDLALEAKSLISKIPGISVLEFPGFSSFFHIDPLRFTIGVWQLGLSGFEADDILCKDFG<br>VVCELVGTKSFTLAFNLGTQRDHVLRLVAGVKHLSKTSHLHQPLKDEVKNVNHFACFDDVRMSMSP<br>REAFFAEKRKVSIRDSLGEICGELVCPYPPGIPVLIPGEIITEKALSYLLEIRSKGAVITGAADCSLSSFLV<br>CVT |
| *Nicotiana tabacum* | NtOLADO A0A1S3ZKS9 | UniProt / Sierro *et al* 2014 genome[3] | ELPPLVNAIKISVEKNATPFHFPGHKRGVAAPSSLVDIISIAPFLYDATELPELDKFGYPTGPLLDAQNQ<br>AAELFGATETWFLVCGTTCGILAAIMSACSPGDTLILARNSHVSATSAMVLCGAIPRYIVPEHNLEWDI<br>AGGVTPSQVKMAIQESEKEGKRAAAVFVTSPTYNGVCSNLSEISQICHFHGIPLIVDEAHGAHFKFHP<br>NMPKTALSQGADLVIQSTHKVLCSFSQSSMLHLSGNRIDRDRVHKCLQSLQTTSPNWLLLASLDATR<br>DELSKNPNTLFNEVMELVQEVKEVIIHIPGISLLDLSSFSNNFSSIDPLRMTIGTQQLGLSGFQAYDILST<br>SHGIEPELIGTKSFTLAVSLGTTKEHSKRLVKGLKYLSTNFLREIKMKRKIIDDNGIEGVPFGEVYMTCT<br>PREAFFARKKIVIFEESIGEVCGEFICPFPPGIPVLIPGEVITKRAVDYLIQVRDQGAFLKGAADPLLASV<br>VVCDF |
| *Artemisia annua* | chr4g02112611 | Liao *et al* 2022 genome[5] | MERTPLVSVLKALADQNVASFHFPGHNRGGAAPSSLSNLIGIQPFLHDLSTVSELDNLFAPAGPILDA<br>QQQAAKLFGATETWFLVGGTTCGIQAAIMATCSPGDTLILPRNAHKSTFSSMVLSGAIPKYITPEYDFD<br>WDLAGSIPPLQVEKAIKELDNQGRKPSAVLVTSPTYHGICSNIEDISILCHSHNIPLIVDEAHGAHLGFH |

| | | | |
|---|---|---|---|
| | | | KNLPRSALSQGADLSIQSTHKVLCSLTQSSMLHMSGNLIQKERICQCLQTLESTSPSYLLLASLDAARAQISENPQTIFNIAVELAFEAKALIEKIPGITVLNSDDPLRITVGLWKLGISGFEANHILKKIGVVAELPGTRSVTFAMNLGTSRDDVKRLVLGLEHLSQMHSLIRGNEEERNDLRAFMETINGMRLSPREAFFASKMKVSFKESIGRICGELVCPYPPGIPLLIPGEVITKEALSYLTGILNKGGFVIGVADPSLSSIIVC |
| *Artemisia annua* | chr7g04669011 | Liao *et al* 2022 genome[5] | MNHTPLVTALKASAKQNVASFHFPGHNRGRAAPSSISNLIGIQPFLHDLPELPELDNLFSPEGPILDAQEQAAKLFGAAETWFLVGGTTCGIQAAIMATCSPGDTLILPRNAHISAFSSMVLSGVIPKYIIPEYDFDWDIACGVTPSQVENAIKESEIEGRKASAVLITSPTYHGICSNLEEISLLCHSHNIPLIVDEAHGAHLGFHKNLPRSALSQGVDISVQSTHKVLCSLTQSSMLHISGNIVNRERICQCLQTLQSTSPSYLLLASLDAARAQISENPKTIFSKAIEIAIEAKYLIEKIPGITVLNSDDPLRVTVGVWNLGISGFEADDILYESYGVVSELTGTRSITFAINLGTSRDDVLRLVSGLQHLSQTYKLVPLKEERLNDIQSFMETSSGMRLSPREAFFASKRKVTFKESIGNVCGELVCPYPPGIPILIPGEVITEEALGYLVDVKKNGGFVSGAADPSLSSIVICT |
| *Artemisia annua* | chr7g04669261 | Liao *et al* 2022 genome[5] | MHDPPLVKTLKTLAEQNVANFQFPGHNRGRAAPSSLSEVIGIQPFLHDMHALPELGNLFSREGAIWDAQKQAAELFGASETWFLVGGTTCGIQASIMATCSPGDTLILPRNAHKSTFSSMVLCGAIPKYLFPEYDYEWDIAGGITPSQVEKAIKELDTEGRKASAVLITSPTYHGICSNLEKISFLCHSYNIPLIVDEAHGAHLGFHENLPRSALSQGVDLSIQSTHKVLCSFTQSSMLHMSGNIVDRERISQCLQSLQTTSPNHMLLASLDAARAQISKHPKTIFNKPFELAMDAKALIEKIPGITVLNSCTFSDVVGIDSLRITVGVWKLGISGFEADCILYKNGVISELQDARSLMFAINLGTTRDDVVKLVEGLKNLSQRHASIRVNVVQPFIGTVSNMILSPREAFFTSKMKVSLKGSIGKVCGELVCPFPPGIPLLVPGEVITAEALNYIVEVKNNGGFISGVADSTLSSIVVCIQ |
| *Artemisia annua* | chr7g04669441 | Liao *et al* 2022 genome[5] | MTYPPLVKTLKALAEQNVANFQFPGHNRGQAAPSSLSGVIGIEPFLHDMHALPELGNLFSREGAISDAQKQAAELFGASETWFLVGGTTCGIQASIMATCSPGDTLILPRNAHKSTFSSMVFCGSIPKYIFPEYDYEWDIAGGITPSQVEKAIKDLDSEGRKASAVLITSPTYHGICSNLEKISFLCHSHNIPLIVDEAHGAHLGFHENLPRSALSQGVDISIQSTHKVLCSFTQSSMLHMSGNIVDRERISQCLQSLQTTSPNHMLLASLDATRAQISENPKTIFNKPFELAMDAKALIEKIPGITVLNSCTFSDVVGLDSLRITVGVWKLGISGFEADRILYKNGVISELQGARSLMFAINLGTSRDDVVKLVSGLKYLSQKHASIQVHDIKPFIGTVSNMILSPREAFFASKTKVGLKESIGKVCGELVCPFPPGIPLLIPGEVITKEALSYLVEVKRNGGFVTGIADSSLRSIVVCG |

**Table S4. Primers used for mutagenesis.** Designed for to mutate the codon optimised sequences on a pET28a plasmid.

| Primer name | Substitutions | Parent | Forward | Reverse |
|---|---|---|---|---|
| OLAD-E117G | E117G | OLAD-WT | CTTCCCG**GT**TTACCTGAACTGGATAATC | GTAA**AC**CGGGAAGGTCATGCAGAAACG |
| OLADO-G170E | G170E | OLADO-WT | CTGACCG**AA**CTTCCAGATATTGACGCCTTG | GGAAG**TT**CGGTCAGGTCATACATAAACG |
| OLAD-P116T | P116T | OLAD WT | CTT**A**CCGAATTACCTGAACTGGATAATC | GTAATTCGG**T**AAGGTCATGCAGAAACG |
| OLADO-T169P | T169P | OLADO WT | CTG**C**CCGGTCTTCCAGATATTGACGCCTTG | GGAAGACCGG**G**CAGGTCATACATAAACG |
| OLAD-P116T+E117G | P116T+E117G | OLAD-WT | CTT**A**CCG**GT**TTACCTGAACTGGATAATC | GTAA**AC**CGG**T**AAGGTCATGCAGAAACG |
| OLADO-T169P+G170E | T169P+G170E | OLADO-WT | CTG**C**CCG**AA**CTTCCAGATATTGACGCCTTG | GGAAG**TT**CGG**G**CAGGTCATACATAAACG |
| OLAD_Y331N | Y331N | OLAD WT | CCCAGC**A**ATTTACTGCTTGCGAG | GTAAA**T**TGCTGGGAGAGGTCGACTG |
| OLADO_N384Y | N384Y | OLADO WT | CCAAT**T**ATTTATTGTTGGCCTCTC | CAATAAAT**A**ATTGGCACTTGTTGACTGC |
| OLAD_Y330N | Y330N | OLAD WT | CCCA**AT**TATTTACTGCTTGCGAG | GTAAATA**AT**TGGGAGAGGTCGACTG |
| OLADO_N383S | N383S | OLADO WT | CCA**GC**AATTTATTGTTGGCCTCTC | CAATAAATT**GC**TGGCACTTGTTGACTGC |
| OLAD_S330N+Y331N | S330N+Y331N | OLAD WT | CCCA**ATA**ATTTACTGCTTGCGAG | GTAAAT**TAT**TGGGAGAGGTCGACTG |
| OLADO_N383S+N384Y | N383S+N384Y | OLADO WT | CCA**GCT**ATTTATTGTTGGCCTCTC | CAATAAAT**AGC**TGGCACTTGTTGACTGC |
| OLAD_Y507F | Y507F | OLAD WT | CCGT**T**CCCCCCAGGCATACC | GGG**A**ACGGGCACACCAGTTCG |
| OLADO_F560Y | F560Y | OLADO WT | CCTT**A**CCCTCCAGGCATACCAG | GGG**T**AAGGACAGATTAATTCC |

**Table S5. OLADO correlation *N. benthamiana*.** All-by-all pearson correlation coefficient (PCC) calculated by Plant Expression Omnibus[4], across 476 samples. Selection of genes with top 30 PCC values with Scf02530g05010 (OLADO) query. UniProt accession values for queries: PMT1_TOBAC (PMT1, putrescine methyltransferase), AO2A_TOBAC (AO2, aspartate oxidase), IFRH_TOBAC (A622), QPT2B_TOBAC (QPT2, quinolinate phophoribosyltransferase), QS1_TOBAC (QS1, quinolinate synthase), F1T160_TOBAC (BBL, berberine bridge enzyme), and MPO1_TOBAC (MPO1, N-methylputrescine oxidase). Pearson correlation coefficient of ODC orthologs (Scf01580G05004 and Scf03865G03002 (top blast hits with ODC1B_TOBAC) with Scf02530g05010 added subsequently.

| Gene | Annotation | PCC |
|---|---|---|
| NIBEN101CTG12894G00002.1 | OLADO (fragment) | 0.958 |
| NIBEN101SCF06783G00003.1 | PMT1 | 0.931 |
| NIBEN101SCF03006G02002.1 | AO2 | 0.93 |
| NIBEN101SCF00918G00001.1 | AO2 | 0.921 |
| NIBEN101CTG08518G00003.1 | PMT1 | 0.921 |
| NIBEN101CTG08674G00002.1 | BBL | 0.913 |
| NIBEN101SCF03045G02019.1 | A622 | 0.9 |
| NIBEN101SCF03045G01004.1 | | 0.899 |
| NIBEN101SCF03687G01012.1 | QPT2 | 0.897 |
| NIBEN101SCF00293G07014.1 | BBL | 0.89 |
| NIBEN101SCF02552G00006.1 | | 0.881 |
| NIBEN101SCF03045G02020.1 | | 0.881 |
| NIBEN101SCF01112G02010.1 | | 0.876 |
| NIBEN101SCF06357G03002.1 | BBL | 0.876 |
| NIBEN101SCF04788G00023.1 | QPT2 | 0.86 |
| NIBEN101SCF00039G01003.1 | BBL | 0.859 |
| NIBEN101SCF07320G00016.1 | | 0.856 |
| NIBEN101SCF02077G13016.1 | MPO1 | 0.852 |
| NIBEN101SCF03674G01015.1 | | 0.848 |
| NIBEN101CTG11222G00002.1 | | 0.844 |
| NIBEN101SCF09960G01015.1 | QS1 | 0.826 |
| NIBEN101SCF08195G07026.1 | | 0.82 |
| NIBEN101SCF15645G02001.1 | MPO1 | 0.819 |
| NIBEN101SCF00069G03001.1 | | 0.809 |
| NIBEN101SCF02049G00002.1 | | 0.794 |
| NIBEN101SCF03029G03002.1 | | 0.792 |
| NIBEN101SCF05437G04016.1 | QS1 | 0.788 |
| NIBEN101SCF01574G00002.1 | | 0.785 |
| NIBEN101SCF05918G03001.1 | | 0.784 |
| NIBEN101SCF05469G03004.1 | | 0.776 |
| NIBEN101SCF01580G05004.1 | ODC | 0.474 |
| NIBEN101SCF03865G03002.1 | ODC | 0.442 |
